# Oxidative Modifications of Parkin Underlie its Selective Neuroprotection in Adult Human Brain

**DOI:** 10.1101/2020.02.19.953034

**Authors:** Jacqueline M. Tokarew, Daniel N. El-Kodsi, Nathalie A. Lengacher, Travis K. Fehr, Angela P. Nguyen, Brian O’Nuallain, Ming Jin, Jasmine M. Khan, Andy C. H. Ng, Juan Li, Qiubo Jiang, Mei Zhang, Liqun Wang, Rajib Sengupta, Kathryn R. Barber, An Tran, Stephanie Zandee, Xiajun Dong, Clemens R. Scherzer, Alexandre Prat, Eve Tsai, Masashi Takanashi, Nobutaka Hattori, Jennifer A. Chan, Andrew B. West, Arne Holmgren, Lawrence Puente, Gary S. Shaw, Gergely Toth, John M. Woulfe, Peggy Taylor, Julianna J. Tomlinson, Michael G. Schlossmacher

## Abstract

The mechanisms by which Parkinson disease-linked parkin confers neuroprotection of human dopamine cells remain elusive. We hypothesized that its cysteines mediate multiple anti-oxidant effects in the midbrain. By studying >60 control specimens, we found that in adult human brain - but not in skeletal muscle- parkin is mostly aggregated and insoluble due to oxidative modifications, such as at C253. *In vitro*, parkin’s oxidation directly reduces hydrogen peroxide (H_2_O_2_) to water. In parkin-deficient human brain, H_2_O_2_ concentrations are elevated. In dopamine toxicity studies, wild-type parkin -but not disease-associated mutants-prevents neural death by lowering H_2_O_2_ and sequestering radicals within insoluble aggregates. Parkin conjugates dopamine metabolites at the human-specific residue C95 and augments melanin formation *in vitro*. Using epitope-mapped antibodies, we found that in adult *Substantia nigra* neurons parkin localizes to neuromelanin within LAMP-3/CD63-positive lysosomes. We conclude that parkin’s own oxidation, previously considered a loss-of-function event, underlies three neuroprotective effects in adult midbrain: its cysteines participate in H_2_O_2_ reduction, dopamine radical conjugation and the formation of neuromelanin.

## INTRODUCTION

Parkinson disease (PD) remains an incurable disorder of the human brain. Bi-allelic mutations in the *PRKN* gene, which encodes parkin, lead to young-onset, autosomal-recessive PD (ARPD)^1^. Clinicopathological studies of parkin-deficient brains have demonstrated restricted cell loss in the *S. nigra* and *L. coeruleus*, two nuclei synthesizing dopamine (DA). The field has not yet explained the selective neurodegeneration of *PRKN*-linked ARPD *vs.* other forms of PD.

Parkin, a principally cytosolic protein, has been associated with diverse cellular functions, foremost related to ubiquitin ligase activity and mitochondrial integrity^2^. However, none of these roles has explained its neuroprotective selectivity, and animal models of genomic *prkn* deletion have failed to reproduce DA cell loss and parkinsonism. This observation could be due to compensatory mechanisms, a shorter life span, or the uniqueness of human DA metabolism. The latter is exemplified by the generation of neuromelanin in dopaminergic cells beginning in late childhood^3,4^. Nevertheless, select *prkn*-null models have revealed changes in high energy-producing cells of flies^5^ and murine brain indicative of augmented oxidative stress^6-8^. We postulated that wild-type parkin promotes redox homeostasis *in vivo.*

Redox equilibrium frequently involves cysteine-based chemistry; there, thiols are subject to oxidative modifications by reactive oxygen-, reactive nitrogen- and reactive electrophilic species (ROS, RNS, RES)^9^, some of which are reversible. Proteins irreversibly conjugated by RES, including by select DA radicals, are degraded, secreted or sequestered within inclusions. It is thought that this process underlies neuromelanin formation^10^.

Human parkin contains 35 cysteines^1^. Several reports previously demonstrated its sensitivity to ROS and RES in cells^11,12^. RNS also attack parkin, and NO-/NO_2_-modified variants have been described in cells and tissue isolates^13^. Such oxidative modifications, which reduce parkin’s ubiquitin ligase activity *ex vivo*, have been previously interpreted as a ‘loss-of-function’^12,14-16^ *in vivo*. In contrast, we hypothesized that parkin’s oxidation ensures redox homeostasis by actively neutralizing ROS, RNS and DA radicals in human brain, and that in parkin’s absence their accumulation leads to selective neurodegeneration in *PRKN*-linked ARPD. Hence, we explored parkin’s metabolism in adult human midbrain and pons *vs.* cortex, spinal cord, skeletal muscle, tested its contribution to redox equilibrium, and studied parkin modifications during DA metabolism.

## RESULTS

### Parkin is mostly insoluble in the ageing human midbrain

Parkin’s metabolism in the human nervous system has remained largely unexplored^17^. We serially fractionated > 40 cortices (age range, 5-85 years) and 20 midbrain specimens (26-82 years) from individuals who had died from a non-neurological cause (**Fig. 1, Extended Data Fig. 1; Supplementary Table 1**). We found that before age 20 years, nearly 50% of cortical parkin was found in soluble fractions generated by salt [Tris-NaCl]- and mild detergent [Triton X-100]-containing buffers (**Fig. 1a,b; Extended Data Fig.1a**). In contrast, after age 50 years, parkin was found nearly exclusively (> 90%) in the 2% SDS-soluble fraction and 30% SDS extract of the final pellet in brainstem and forebrain specimens, which included the *S.nigra, L.coeruleus*, striatum and cortex (**Fig. 1a-c**). The requirement for SDS to solubilize parkin suggested it had been aggregated, which is consistent with its co-distribution with LC3B, a phagosomal marker of aggregated proteins (**Fig. 1d**). In soluble brain fractions from older humans, we did not detect any truncated species of parkin using several, specific antibodies. Despite the protein’s solubility loss, *PRKN* mRNA was detectable in individually examined nigral and cortical neurons from all age groups **(Fig. 1e**). Moreover, transcript levels were independent of age (**Extended Data Fig. 1d**).

**Figure 1:**
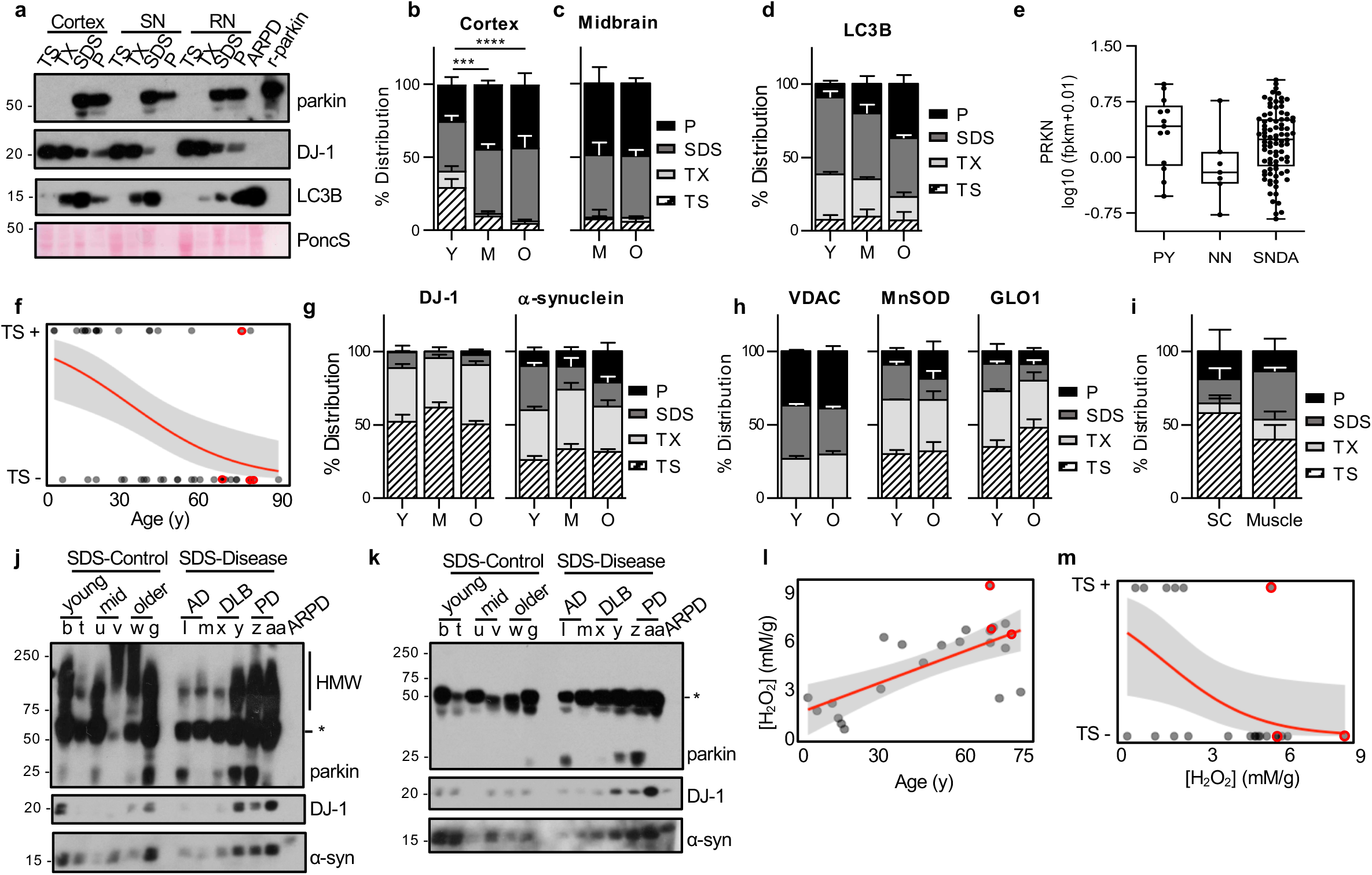
Parkin transitions from a soluble to an aggregated state in adult human midbrain. **(a)** Representative Western blots of parkin, DJ-1, and LC3B distribution in human cortex, *S. nigra* (SN) and red nucleus (RN) brain specimens that had been serially fractionated into Tris-NaCl buffer-soluble (TS), Triton X-100-soluble (TX), 2% SDS-soluble (SDS) extracts and the pellet (P) lysed in 30% SDS-containing buffer. Lysates from *PRKN*-linked Parkinson disease (ARPD) brain and recombinant, human parkin (r-parkin) are included. **(b-c)** Relative distribution of parkin signal within each fraction for **(b)** cortex and **(c)** midbrain grouped by age ranges: young (Y; ≤ 20y; n=13); mid (M, > 20y, < 50y; n=15 cortex, 6 midbrain); older (O, > 50y; n=13 cortex, 14 midbrain). Data shown as mean ± SEM. Significance in protein distribution between soluble (TS+TX) and insoluble (SDS+pellet) fractions was determined using 2-way ANOVA (***p < 0.001; ****p < 0.0001). Additional Western blots in **Extended Data Fig. 1a-c**. **(d)** Signal distribution for LC3B in human cortices as in **(b)**. (n=3-5 per age group). **(e)** Quantification of log-transformed *PRKN* mRNA signals from individual pyramidal neurons (PY), non-neuronal leukocytes (NN) and *S. nigra* dopamine neurons (SNDA) isolated from healthy controls (ages, 59-89y). **(f)** Logistic regression analysis of parkin solubility in cortices as a function of age (n=45). Each brain is represented by an individual dot; red dots denote cases of late-onset Parkinson’s not linked to *PRKN*; the logistic regression line (in red) and 95% confidence intervals (grey) are shown. **(g-h)** Relative distribution of DJ-1, α-synuclein **(g)** and **(h)** VDAC, MnSOD and GLO1 as described in (b). Representative Western blots in **Extended Data Fig. 1b,c**. **(i)** Relative distribution of parkin as in (b) for human spinal cord (n=4) and skeletal muscle specimens (n=6), from donors aged 50-71y. Representative Western blots are in **Extended Data Fig. 1e.** **(j-k)** Western blots of parkin, DJ-1, and α-synuclein in SDS cortical fractions from healthy controls (3 age groups) and 6 patients including Parkinson’s (PD), Alzheimer disease (AD) and dementia with Lewy bodies (DLB). The same lysates were separated by SDS-PAGE under **(j)** non-reducing and **(k)** reducing conditions. **(l-m)** Linear regression analyses of **(l**) H_2_O_2_ concentrations in control cortices (mM/g tissue) as a function of age for control cortices, and **(m)** logistic regression analyses of parkin solubility as a function of H_2_O_2_ levels in the same specimens (n=20) as in (f). Red circles denote three disease cortices (AD; DLB; PD).

Using a logistic regression model with parkin as the dependent variable, we found that its solubility in human brain correlated with age (**Fig. 1f**), where the age coefficient was −0.0601 (95% CI: −0.106 to −0.024; P=0.004). When using age to classify soluble *vs.* insoluble, the area under the receiver operating characteristic (ROC) curve was 0.8. This transition to insolubility occurred between the age range of 28 (low sensitivity; high specificity) and 42 years (high sensitivity; low specificity). We confirmed that parkin’s solubility change was not caused by tissue freezing, was independent upon *post mortem* interval (PMI) and sex, and it was not affected by the pH of extraction buffers (**Extended Data Fig. 2**; not shown).

The age-dependent partitioning of parkin was not seen for any other protein examined, which included PD-linked proteins, *e.g.*, DJ-1 and α-synuclein (**Fig. 1a,g)** and organelle-associated markers, *e.g.:* cytosolic glyoxalase-1, peroxiredoxin-1 and −3; endoplasmic reticulum-associated calnexin; and mitochondrial VDAC and MnSOD (**Fig. 1h; Extended Data Fig. 1b,c;** and data not shown). Interestingly, its partitioning was organ- and species-specific. In contrast to human brain, parkin remained soluble in skeletal muscle and spinal cord specimens from older individuals (**Fig. 1i; Extended Data Fig. 1e**). In the brains of aged mice (at 22 months), rats (14 months) and cynomolgus monkey (5 years), parkin remained soluble even when the length and conditions of PMIs were matched to those of human tissues (**Extended Data Fig. 1f**; data not shown). We concluded that parkin’s solubility loss is relatively unique to human brain ageing.

### Parkin is oxidized in adult human brain

When human brain fractions were examined under non-reducing conditions, we detected parkin proteins ranging in *M*_*r*_ from > 52 to 270 kDa, invariably in the form of smears (**Fig. 1j**). These species were abundant in SDS extracts and reduced to parkin’s monomer (∼52.5 kDa) by adding dithiotreitol (DTT) or β-mercaptoethanol (β-ME). This is shown, for example, in specimens from both normal and neurologically diseased cases (**Fig. 1j,k**). No such reactivity was seen in SDS-extracts of *PRKN-*linked ARPD brains, demonstrating specificity. Examples of oxidative modifications are shown in **Extended Data Fig. 3.** Under the same conditions, high *M*_*r*_ protein variants were rarely seen for other PD proteins, *e.g.*, DJ-1 and α-synuclein. The latter showed oligomers only in select brains from older subjects (not shown). Of note, the formation of high *M*_*r*_ parkin was not due to oxidation *in vitro*, as brain specimens were fractionated in the presence of iodoacetamide (IAA) to protect thiols; they also did not arise from ubiquitin adducts, which cannot be removed by reducing agents.

These findings demonstrated that a pool of parkin had undergone oxidation *in vivo*. Indeed, using logistic regression analyses, we found that in the same brain specimens H_2_O_2_ concentrations significantly correlated with age (**Fig. 1l**). Moreover, parkin’s solubility loss paralleled the rise in H_2_O_2_; the latter’s coefficient was −0.939 (95% CI: −2.256 to −0.248; P=0.0415; **Fig. 1m**).

### Redox stressors directly modify parkin and alter its solubility

To investigate mechanisms by which its solubility is altered by redox stress, we exposed recombinant (r-), tag-less, human parkin to oxidants *in vitro*. The exposure of highly purified r-parkin to rising concentrations of H_2_O_2_, DA, the DA metabolite aminochrome, and mitochondrial toxins, *e.g.*, CCCP and rotenone, generated high *M*_*r*_ smears and progressive insolubility (**Fig. 2a; Extended Data Fig. 4a-c)**). Some of these biochemical changes were reversible, as seen for insoluble, H_2_O_2_-oxidized r-parkin upon exposure to DTT (**Fig. 2b**). Others could not be reduced, such as when irreversible, covalent modifications had occurred (**Extended Data Fig. 4a-c**; data not shown). Blocking access to parkin’s thiols prevented its oxidation (see below). Demonstrating sensitivity to bi-directional redox changes, exposure of native r-parkin to excess DTT also rendered it insoluble (**Extended Data Figs. 3,4d**).

**Figure 2:**
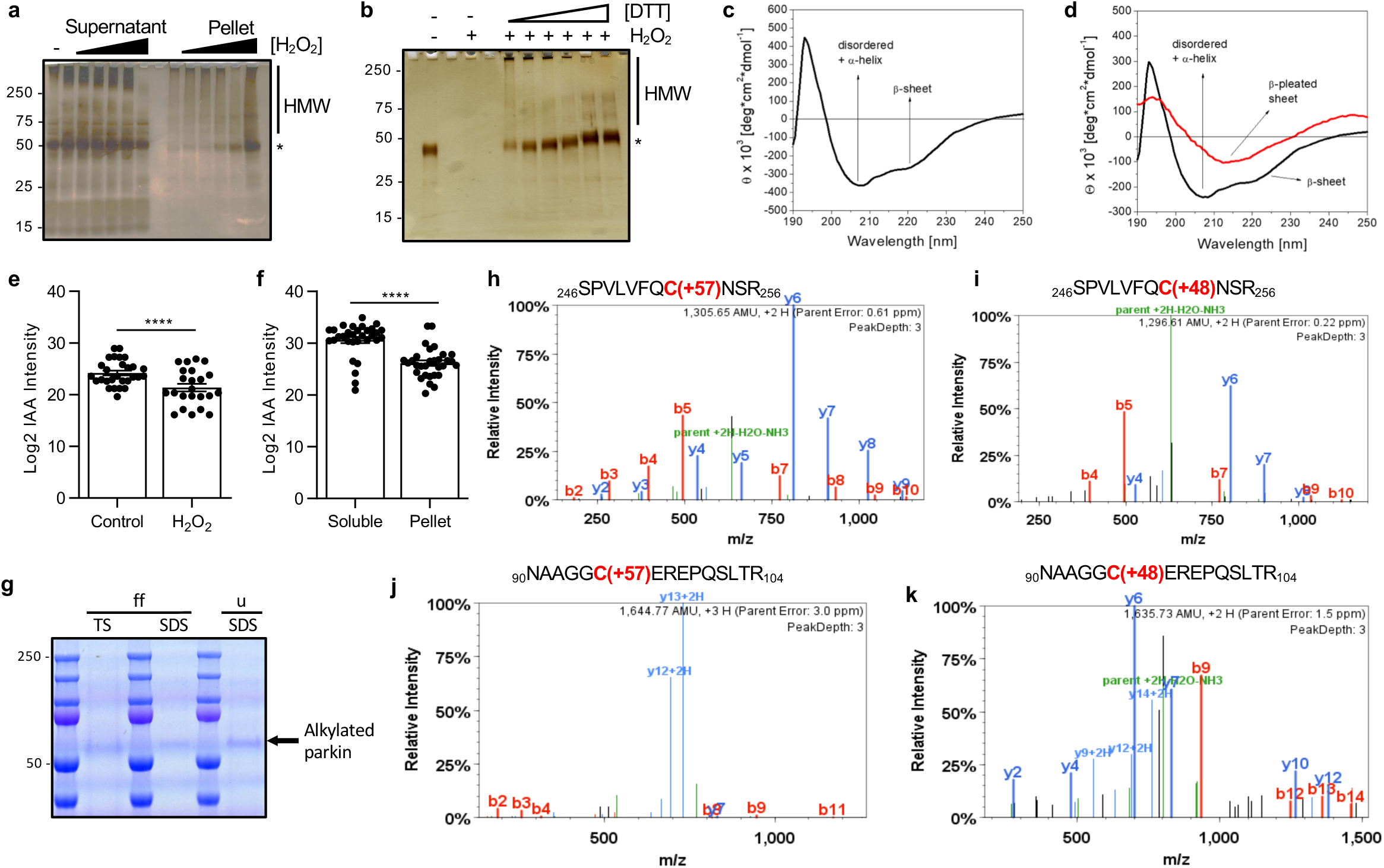
Parkin’s secondary structure is altered by redox stress. **(a)** Silver staining of r-parkin in soluble (supernatant) and insoluble (pellet) phases following exposure to increasing concentrations of H_2_O_2_ (0-2mM) and run under non-reducing conditions. Monomer (*) and high *M*_*r*_ weight (HMW) r-parkin species are indicated. **(b)** Silver stained gel of r-parkin exposed to H_2_O_2_ (10 mM), followed by treatment with increasing concentrations of DTT (0-100 mM) prior to centrifugation and loading of the supernatant onto SDS-PAGE. **(c,d)** Circular dichroism spectra of soluble, untreated r-parkin at **(c)** T=0 and **(d)** soluble (black line) and aggregated (red line) states following incubation at 37°C for T=5 days. The protein secondary structure shifts from a predominant appearance of α-helix dominated state, as demonstrated by the positive band at 193 nm and negative bands 208 nm (black lines), to the appearance of β-pleated sheet formation, as demonstrated by negative bands at 218 nm and a rise in molar ellipticity with positive bands at 195 nm (red lines) during spontaneous oxidation. **(e-f)** Quantitative analyses of IAA-modified cysteines captured by LC-MS/MS for **(e)** untreated *vs.* H_2_O_2_-exposed r-parkin, and **(f)** soluble compared to insoluble (pellet) fractions. Each data point represents the log2-transformed total IAA-signal intensities of single cysteine residues (n=3 runs for each). The cysteine pool is shown with the mean ± SEM; significance **p < 0.01 as determined using Student T-Test. **(g-k)** LC-MS/MS-generated spectra following trypsin digestion of IAA alkylated WT parkin enriched from human cortex (example shown in (**g)**). **(h-i)** C253 (peptide aa246-256) was detected in both **(h)** a reduced state (IAA labelled; carbamido-methylation, +57 mass shift; peptide score 172.99) and **(i)** an oxidized state (*i.e*, trioxidation, +48 mass shift; peptide score 79.04). **(j-k)** C95 (peptide aa90-104) was also found in both a **(j)** reduced state (IAA labelled; carbamido-methylation, +57 mass shift; peptide score 100.16) and **(k)** oxidized state (trioxidation, +48 mass shift; peptide score 84.47). See also **Extended Data Table 1** (human cortex) and **Extended Data Fig. 4h** (r-parkin).

### Redox stress alters parkin’s structure

The progressive insolubility of parkin with increasing oxidative stress *in vitro* and *in vivo* suggested that the nascent protein responds to redox alterations with structural change(s). We tested this by dynamic light scattering (DLS) and far-UV-circular dichroism (CD). When we monitored r-parkin during spontaneous oxidization using DLS (**Extended Data Fig. 4g**), we observed a shift in r-parkin’s hydrodynamic diameter from 5.1 nm, representing a folded monomer at the start, to multiple peaks with larger diameters 5 hrs later; the latter indicated multimer formation. These oxidation-induced changes were reversible by DTT.

To further monitor structural changes, we analyzed the effects of spontaneous oxidation in naïve and H_2_O_2_-treated r-parkin by CD (**Fig. 2c,d; Extended Data Fig. 4e,f**). At the start, soluble fractions contained both α-helically ordered as well as unstructured proteins (**Fig. 2b; Extended Data Fig. 4e**). After five days, r-parkin preparations were centrifuged, and soluble and insoluble fractions re-analyzed (**Fig. 2d; Extended Data Fig. 4f**). There, we found a marked shift by CD from an α-helically ordered (and unstructured) content to increased amounts of β-pleated sheet-positive r-parkin in insoluble, aggregate-rich fractions. This shift was more pronounced in H_2_O_2_-oxidized protein preparations (**Extended Data Fig. 4f**).

### Hydrogen peroxide modifies parkin at multiple cysteines

To further confirm that oxidation of parkin cysteines led to redox state change and insolubility, we used liquid chromatography-based mass spectrometry (LC-MS/MS). We employed a serial IAA-DTT-NEM fingerprinting approach to differentiate reduced thiols (tagged by IAA) from reversibly oxidized thiols (reduced by DTT and then tagged by N-ethylmaleimide (NEM)). We first asked whether oxidation of r-parkin led to changes in thiol redox state regardless of solubility. At higher H_2_O_2_ concentrations, 32 of 35 cysteines (91.4%) were identified; > 93.8 % were found modified at least once. Using Scaffold PTM-software, we recorded the anticipated rise in NEM-tagged cysteines following H_2_O_2_ treatment (**Extended Data Fig. 4h**). Oxidative modifications were identified at parkin’s RING domains and at non-RING-based cysteines. Total cysteine modifications were quantified by MaxQuant software^18^. As shown in **Fig. 2e**, there was a significant loss of IAA signals for r-parkin’s cysteines in H_2_O_2_-treated samples (P=0.0016), reflecting a decrease in reduced thiols.

In turn, when comparing cysteine oxidation in soluble *vs.* insoluble fractions of both untreated and oxidized r-parkin, IAA signals (*i.e.*, free thiols) were significantly reduced in the pellets (P < 0.0001; **Fig. 2f**). Modifications at other residues, *e.g.*, methionines, did not correlate with r-parkin insolubility (not shown). In these studies, residue C253 (RING1 domain) was frequently detected as oxidized. These results established that H_2_O_2_-induced oxidation of cysteine-based thiols conferred r-parkin’s insolubility.

We next examined whether these thiol oxidations also occur in human brain-derived parkin by LC-MS/MS. Here, we mapped cysteines in reduced and oxidized states for the 52.5 kDa monomer purified from cortex specimens of neurological controls. The latter included irreversible oxidation events, such as to sulfonic acid at C253 (**Fig. 2g-i**) and C95 **(Fig. 2j,k**), as well as reversible ones, *e.g.*, at C253, detected in insoluble parkin (see **Extended Data Table 1**).

### Parkin reduces hydrogen peroxide to water

Redox reactions transfer oxygen, electrons and H^+^ between the reactants. We therefore asked whether parkin’s oxidation led to H_2_O_2_ reduction (**Fig. 3; Extended Data Fig. 5a-f**). Indeed, we found that r-parkin directly lowered H_2_O_2_ levels in a time- and concentration-dependent manner (**Fig. 3a; Extended Data Fig. 5c**). This reaction (*e.g.*, 2R-SH+H_2_O_2_→R-S-S-R+2H_2_O) was dependent on thiol integrity, because pre-treatment with IAA abolished it (**Fig. 3b**). Parkin’s reducing capacity was confirmed in parallel studies employing a different H_2_O_2_ assay^&^.

**Figure 3:**
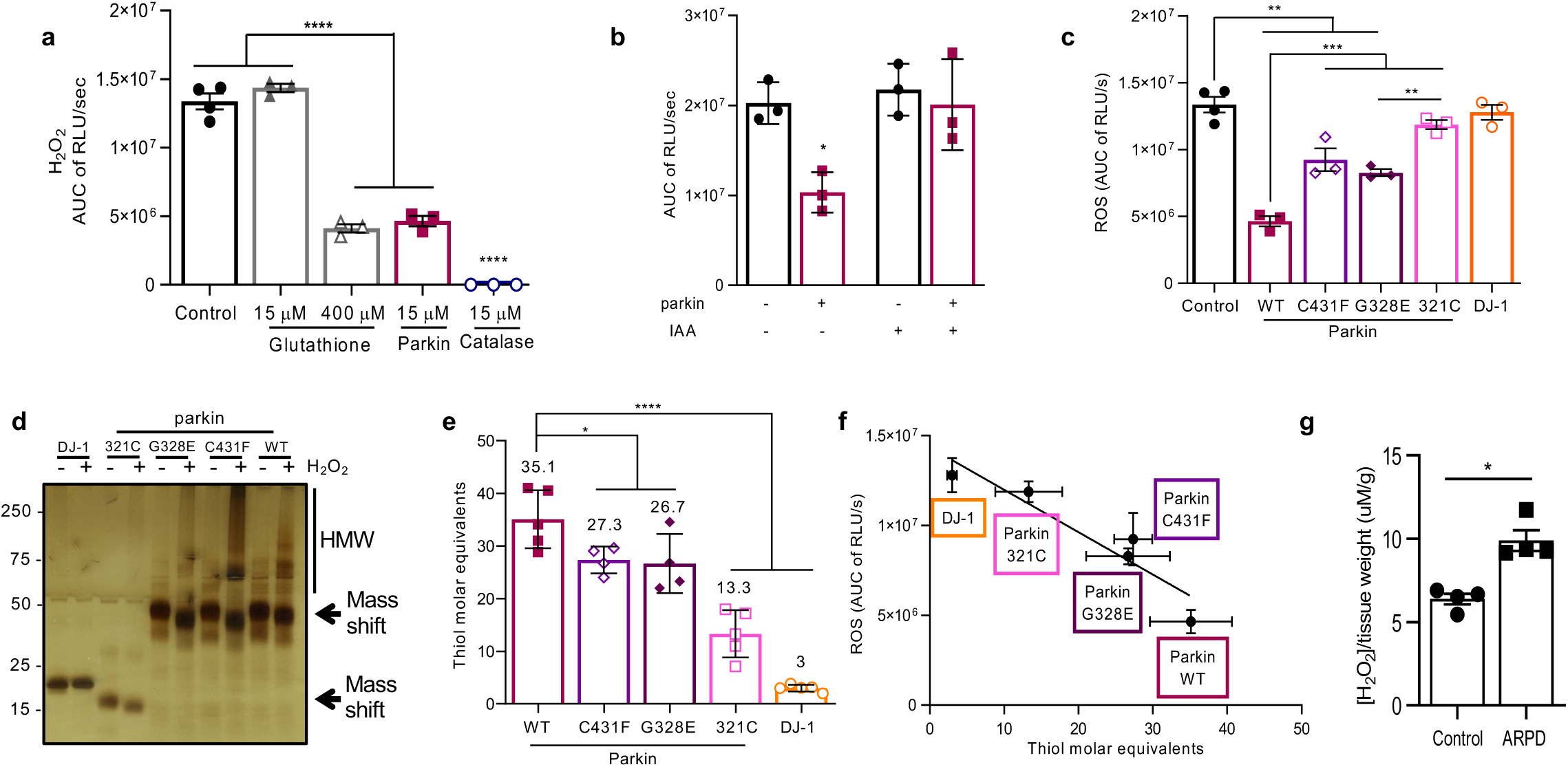
Parkin reduces H_2_O_2_ to water. (**a-d**) Area under the curve (AUC) plots for results from *in vitro* colorimetric H_2_O_2_ assays, where AUC integrates total H_2_O_2_ present over the time course of the assay (see also **Extended Data Fig. 5a**). **(a)** WT full-length r-parkin and glutathione significantly reduced H_2_O_2_; **(b)** This was blocked when r-parkin was pre-incubated with IAA. **(c)** Comparison of WT r-parkin with DJ-1, r-parkin point mutants, and r-parkin_321-465_ (321C). Results represent an average of n=3 ± SD; *p < 0.05, **p < 0.01,***p < 0.001, and ****p < 0.0001 using one-way ANOVA with Tukey’s post hoc test. **(d)** Visualization of proteins by silver staining (non-reducing conditions). **(e)** Quantification of reactive thiol content (in molar equivalents) of r-parkin WT, r-parkin point mutants, r-parkin_321-465_ and full-length r-DJ-1 using the Ellman’s reagent assay. **(f)** Correlation curve between number of free thiols **(e)** and H_2_O_2_ reducing capacity **(c)** for indicated proteins. **(g)** Quantification of H_2_O_2_ in human brain from *PRKN*-deficient ARPD cortices compared to age- and *post-mortem interval*-matched controls collected at the same institution. Results are represented as the mean concentration of H_2_O_2_ per g of tissue analyzed (μM/g; n=4/group) ± SEM; *p < 0.05 determined using Student T-test.

We next examined the reducing capacities of other proteins, *e.g.*, r-DJ-1, a C-terminal RING2 peptide (r-parkin_321C_), and two ARPD-linked mutants, p.G328E and p.C431F (**Fig. 3c-d, Extended Data Fig. 5d-f**). There, r-DJ-1 and r-parkin_321C_ showed negligible H_2_O_2_-reduction, and the point mutants conferred less activity than wild-type r-parkin (**Fig. 3c; Extended Data Fig. 5d,e**). These reducing effects correlated with r-parkin’s own oxidation, as revealed by SDS-PAGE immediately after H_2_O_2_ exposure (**Fig. 3d**). We speculated that this activity by r-parkin was dependent on reactive thiol content. We tested this using the Ellman’s reagent. There, wild-type r-parkin, r-parkin_321C_ and r-DJ-1 showed the predicted number of reactive thiols, whereas the point mutants p.C431F and p.G328E had fewer accessible thiols (**Fig. 3e**). As predicted, we recorded a linear correlation between thiol molar equivalencies and H_2_O_2_-reducing capacities (**Fig. 3f**).

### Hydrogen peroxide is elevated in parkin-deficient brain

To determine whether parkin oxidation conferred ROS reduction in adult brain, we quantified H_2_O_2_ concentration in cortices from human ARPD subjects and age-, PMI-, ethnicity- and brain region-matched control specimens (n=4 each)^1,19^. There, we found a significant elevation in H_2_O_2_ concentrations in parkin-deficient brain lysates (P < 0.05; **Fig. 3g**). Of note, in specimens of non*-PRKN*-linked cases with parkinsonism, H_2_O_2_ levels were similar to those of age-matched controls (Fig. 1l). We concluded that parkin contributes to redox homeostasis in human brain. In related mouse experiments, we identified that parkin’s regulation of neural H_2_O_2_ occurs in the cytosol^&^.

### Parkin prevents dopamine toxicity by lowering hydrogen peroxide

To explore its selective neuroprotection, we revisited the role of parkin in DA-induced toxicity^13^. We first tested its redox-related functions in human neuroblastoma M17 cells. Here, wild-type parkin, but not ARPD mutants p.G328E and p.C341F, protected cells from rising DA levels (**Fig. 4a; Extended Data Fig. 6a**). Moreover, DA-exposure resulted in a significant rise in H_2_O_2_ concentrations, which wild-type-but not mutant-parkin reversed (**Fig. 4b**). These *ex vivo* results validated our *in vitro* and *in vivo* findings (Fig. 3). In stable M17 cells, protection from DA toxicity paralleled *PRKN* expression levels (**Fig. 4c**); approximately 4 ng neutralized each μM of DA. Neuroprotection also correlated with parkin’s own oxidation and progressive insolubility. Importantly, these modifications were not reducible by DTT (**Fig. 4d,e**), indicating irreversible thiol changes, which we reasoned were due to covalent DA adduct formation^13^.

**Figure 4:**
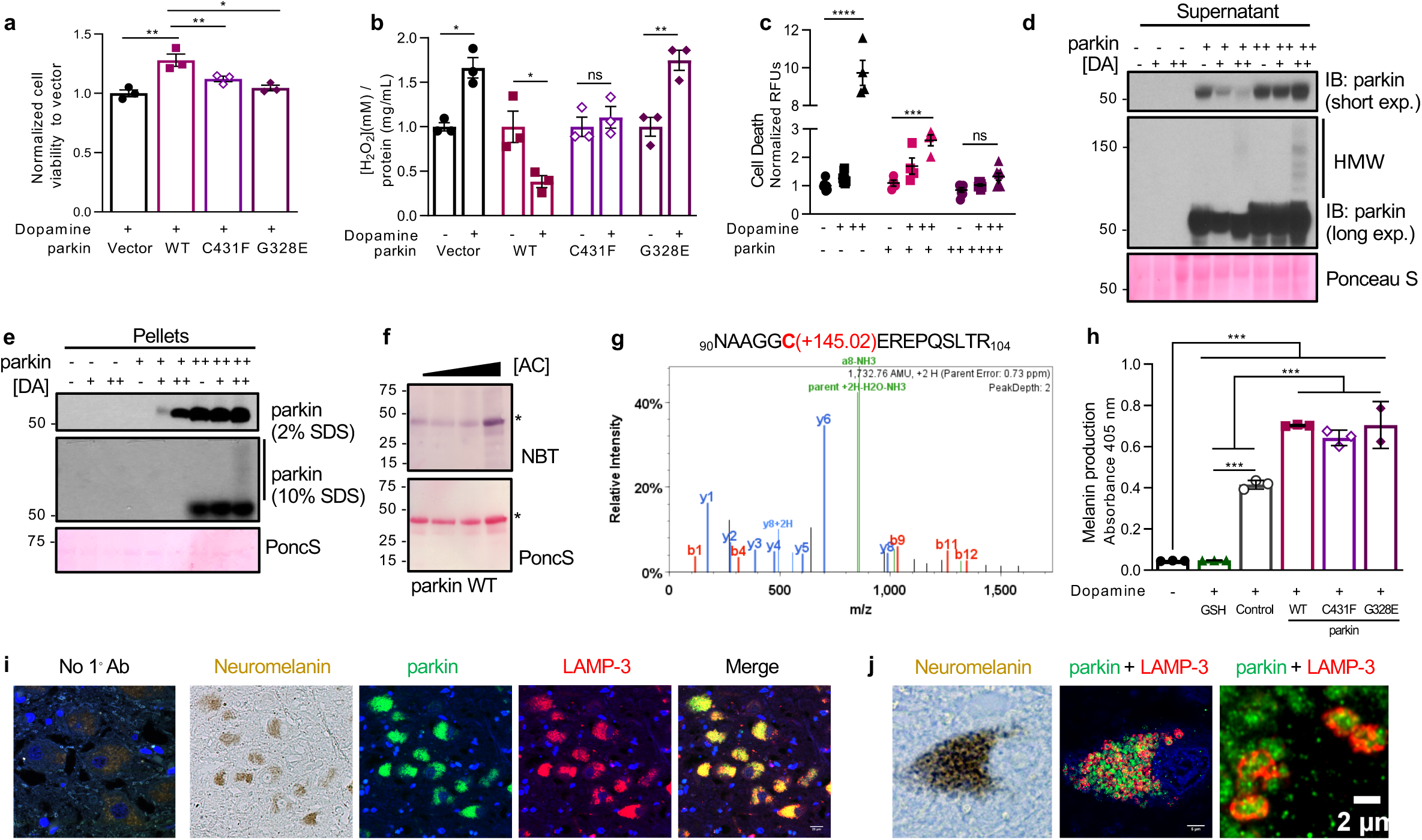
Parkin directly conjugates dopamine metabolites and is associated with neuromelanin *in vivo.* **(a-b)** Cell viability assay **(a)** and **(b)** H_2_O_2_ quantification for dopamine-treated M17 cells expressing WT or two ARPD-linked parkin mutants. Cells were exposed to 200 μM dopamine or vehicle for 20h as indicated. Data represent the mean of duplicates ± SEM. *p < 0.05 by 1-way ANOVA and is representative of n=3. **(c)** Cell viability assay of dopamine-stressed human M17 neuroblastoma cells that stably express vector-control plasmid (parkin -) or human *PRKN* cDNA at mid- (+) or high (++) levels. Cells were stressed with 20 μM (+) and 200 μM (++) dopamine for 20h. Representative data are shown for the mean of duplicates ± SEM from n=4-8 independent experiments (for different concentrations); *p < 0.05 by 1-way ANOVA. **(d-e)** Western blot of parkin in soluble (supernatant, **d**) and insoluble (serial pellets, **e**) fractions of lysates described in **(c)**. SDS/PAGE was run under reducing conditions. **(f)** NBT and Ponceau S staining of WT human r-parkin that had been incubated with increasing concentrations of aminochrome *in vitro*; gel was run under reducing conditions. *Denotes parkin monomers. **(g)** LC-MS/MS-generated spectra highlighting trypsin-digested human r-parkin peptide aa90-104 carrying a indole-5,6-quinone adduct (+145 mass gain) at C95 (Mascot ion score 31.1) See also **Extended Data Fig. 6c,d**. **(h)** AUC of kinetic result of *in vitro* melanin formation as monitored by a colorimetric assay. Dopamine polymerization to melanin was monitored in the presence or absence (blank) of WT, r-parkin point mutants and glutathione (GSH). Data represent the mean of triplicates ± SEM. ***p < 0.05 by 1-way ANOVA. See also **Extended Data Fig. 6b**. **(i-j)** Indirect immunofluorescence microscopy signals generated by anti-parkin (clone-E; green colour) identifies perinuclear organelles throughout the neuronal cytoplasm that are also recognized by anti-LAMP-3/CD63 (red colour), as shown by double-labelling (blue colour, Hoechst stain). Reactivities generated by anti-parkin topographically overlap with juxtanuclear neuromelanin, as shown by panel to the left identifying neuromelanin-carrying cells by light microscopy in the same field. No primary Ab control run in parallel to experiment is shown. Higher magnification of individual dopaminergic neuron is shown in **(j)**.

### Parkin binds dopamine radicals at cysteine 95

We asked whether parkin directly aided in the neutralization of DA through conjugation at specific residues^14^. Indeed, full-length r-parkin bound DA and aminochrome adducts, as detected for example by nitro-blue-tetrazolium staining after aminochrome exposure (**Fig. 4f**; and data not shown). Using LC-MS/MS to map these modifications, we identified several aminochrome, DA and indole-5,6-quinone adduct-carrying peptides specific to treated r-parkin (**Extended Data Fig. 6c,d**). Intriguingly, the most frequently modified residue was C95, a primate sequence-specific cysteine (**Fig. 4g; Extended Data Fig. 6c**).

### Parkin augments melanin formation

Given the dynamic associations between r-parkin, DA exposure, RES conjugation and aggregate formation, we tested whether these influenced melanin formation *in vitro*. Polymerization of DA into melanin occurs in a spontaneous, oxidation-dependent manner via intermediates, such as indole-5,6-quinone, and is blocked by glutathione^10,20^ Wild-type parkin and two point mutants, p.C431F and p.G328E, all augmented melanin formation in a time- and concentration-dependent manner (**Fig. 4h; Extended Data Fig. 6b**). We next sought to validate a role for parkin in melanin formation *in vivo.*

### Parkin colocalizes with neuromelanin in adult *S. nigra* neurons

To visualize parkin’s subcellular localization in adult brain, we carried out routine microscopy studies using newly developed, monoclonal antibodies to human parkin; these had been epitope-mapped and validated regarding their specificity (*e.g.*, **Extended Data Fig. 7**). When staining serial sections of midbrains from control individuals, three antibodies revealed intense, granular parkin staining in the cytoplasm of pigmented neurons from adults (age > 55 years), albeit at varying intensity levels amongst them (clones-D, -E and -G; **Extended Data Fig.8a**). We counted anti-parkin reactivity in > 83% of anti-tyrosine hydroxylase-positive cells and in many non-pigmented neurons of the midbrain and cortex (**Extended Data Fig. 8b,c**).

By immunohistochemistry, in *S. nigra* neurons anti-parkin signals heavily overlapped with neuromelanin pigment (**Extended Data Fig 8d**). Sections from individuals with parkinsonism (not linked to *PRKN* mutations) showed the same anti-parkin reactivities in surviving neurons and overlapping with extracellular pigment following neuronal degeneration. We found weaker anti-parkin reactivity when examining *S. nigra* sections of younger subjects (≤33 years), which matched their paucity in neuromelanin content (**Extended Data Fig. 8d,e**).

### Parkin localizes to LAMP-3^+^-lysosomes within nigral neurons

In screening organelle markers by immunohistochemistry, we detected highly similar staining between antibodies to parkin and lysosome-associated protein-3 (LAMP-3/CD63) reactivities (**Extended Data Fig. 8f**). Antibodies to LC3B -but no mitochondrial antigens (*e.g.*, MnSOD, VDAC)-generated much less signal overlap (not shown). Because neuromelanin granules are thought to arise within non-degradative lysosomes^21^, we carried out immunofluorescence microscopy. We found that both anti-parkin and anti-LAMP-3 antibodies strongly co-labelled with pigmented granules (**Fig. 4i**). Of note, LAMP homologues have been shown to promote long term storage of melanin in human keratinocytes^22^; moreover, variants at the *LAMP-3* locus have recently been linked to PD susceptibility^23^.

Using confocal microscopy, we demonstrated that in adult midbrain sections anti-parkin signals and neuromelanin granules were frequently surrounded by anti-LAMP-3 reactivity (**Fig. 4j**). We concluded that parkin metabolism is likely associated with LAMP-3-mediated-sequestration of DA radicals in the *S. nigra* of human midbrain.

## DISCUSSION

We identified a heretofore unrecognized role for parkin in redox homeostasis of human brain. To date, only one other protein reportedly has a similar duality in functions (*i.e.*, ubiquitylation and redox biology), namely the sensitive-to-apoptosis gene product, SAG (aka RBX2/ROC2/RNF7)^24^. Our work establishes that parkin’s progressive insolubility in human brain, first described in the cortex by Pawlyk et al.^17^, is due to oxidative modifications during normal ageing (**Fig. 1, 2d-k; Extended Data Table 1**). Unlike in adult brain, where parkin is mostly insoluble after the fourth decade, in the human spinal cord and aged skeletal muscle ∼50% of detectable parkin remains soluble. In all other aged mammals we examined, brain parkin remains soluble to an even greater degree (**Fig. 1i; Extended Data Fig. 1.e,f**). Theoretically, other modifications (*e.g.*, phosphorylation^25^) could also alter parkin’s solubility.

Heretofore, parkin oxidation in cells has been interpreted as a ‘loss-of-its-function’ *in vivo*. In contrast, we posit that parkin’s post-translational oxidation, related aggregate formation, and translocation into lysosomes all reflect its physiological function to maintain redox balance. Ubiquitylation and redox stabilization may not be mutually exclusive. Low concentrations of pro-oxidants have been shown to initiate parkin’s ubiquitin ligase activity *in vitro*^15^, which we confirmed^&^.

The cellular thiol network is critical in DA cell health^26^. As summarized in **Extended Data Fig. 9**, through its oxidation we postulate that parkin neutralizes ROS (*e.g.*, H_2_O_2_ to water; **Fig. 3**), reduces RNS, conjugates RES (*e.g.*, quinones), promotes melanin formation, and sequesters radicals within insoluble aggregates (**Fig. 4**). This multimodal redox function is supported three-fold. One, in mice by evidence of sustained oxidative stress in parkin’s absence^6,7^. Two, in humans by complementary findings in ARPD brain (**Fig. 3**) and the observation of lesser neuromelanin content in surviving DA neurons^27-30,51-52^. We noted with interest that primate-specific residue C95 (located outside of its RING domains) was most frequently found to conjugate DA radicals (**Fig. 4g; Extended Data Fig. 6c**). And three, in parallel work, we discovered a dynamic feedback loop between *PRKN* expression and glutathione levels *in vivo*^&^.

We found that single point mutants have measurable redox effects depending on the experimental context explored (**Fig. 3, 4**). Such complementary studies will inform clinical genotype-to-phenotype studies in humans^31,32^. Several reports have shown that point mutation carriers have a later age-of-onset of ARPD than those whose alleles encode no RING domain^31,32^. The subcellular translocation of parkin (to mitochondria or lysosomes) could be facilitated by oxidation-induced multimer formation at its RING domains (**Fig. 2g,h**). This concept of ‘RING scaffolding’ has previously been described for other ubiquitin ligases^33^. Because neuronal health and axonal integrity are co-regulated by redox chemistry^34^, we posit that parkin confers its ‘inherently present, pro-mitochondrial benefit’^35^ via redox homeostasis.

Our study raises new questions. Beyond its localization within LAMP-3^+^-lysosomes in DA cells (**Fig. 4i,j**), our results have not yet revealed the fate of oxidized parkin in other neurons. Human LAMP proteins have been shown to store melanin in human keratinocytes to protect them from UV light^22^. Thus, oxidized parkin together with LAMP-3 (and possibly, LAMP-2) could play a similar role in the ongoing storage of neuromelanin and other radicals, thus facilitating their effective sequestration in ageing brain. Of note, the three clones that detect parkin by microscopy seem to preferentially bind to oxidized/aggregated forms *in situ*. Little anti-parkin signal was detected in midbrain sections from young adults (**Extended Data Fig. 8d**), whereas the holoprotein was readily detected by immunoblotting (*e.g.*, **Fig. 1a**).

Several limitations were identified. One, although we have detected the *PRKN* gene to be continuously transcribed in nigral neurons, even at age 99 years (**Fig. 1e**), the rate of its mRNA translation under redox stress conditions, such as during ageing, remains to be studied. Future studies will employ sensitive ELISA-based techniques to quantify the precise concentration of nascent parkin in tissues. Two, we have only begun to map parkin’s oxidative modifications that occur in normal human brain (*e.g.*, **Extended Data Table 1**). As highlighted in Extended Data Fig. 3, parkin likely carries a heterogenous complement of modified cysteine residues in human brain, making the completion of qualitative mapping by LC-MS/MS difficult. Moreover, ROS and RNS modifications can be very transient and thus difficult to capture directly^15,16,36^. Three, we have not yet completed a quantitative assessment of oxidative modifications in young *vs.* old human brain extracts. Future MS studies require further optimization and larger amounts of human tissues including from healthy midbrains. In addition, access to midbrain organoids^37^, such as those generated from entirely parkin-deficient patients and subjects carrying *PRKN* point mutants, will provide clarification as to which parkin domains are associated with neuromelanin formation.

The strength of this study is that it focused on parkin metabolism in human brain. *PRKN-*linked ARPD is a human condition, and we are guided by insights from human studies. We posit that parkin fulfils three criteria of an essential redox molecule: the sensing of stressors; structural responses to them; and the participation in corrective measures. If confirmed by future work, this new concept for parkin’s function in the human brain opens the door to revisit a possible role in non-*PRKN*-linked PD and in parkinsonism associated with advanced ageing. It also allows the revisiting of therapeutic approaches in patients afflicted by ARPD, such as through viral vector delivery of *PRKN* cDNA into parkin-deficient brain^38^ and/or by polyvalent, anti-oxidant therapy in select patients^39^.

## MATERIALS AND METHODS

### Tissue collection

All tissue was collected in accordance with Institutional Review Board-approved guidelines. Fresh frozen samples of cortical human brain from subjects under 50 years of age were acquired through the University of Alabama and the Autism Tissue Program. *Post mortem*, frozen brain samples from frontal cortices were also obtained from the NICHD Brain and Tissue Bank at the University of Maryland. Brain tissues, including midbrain specimens, with short PMI were also obtained from patients diagnosed with clinical and neuropathological MS according to the revised 2010 McDonald’s criteria (n=4) ^40^. Tissue samples were collected from MS patients with full ethical approval (BH07.001) and informed consent as approved by the CRCHUM research ethics committee. Autopsy samples were preserved and lesions classified using Luxol Fast Blue / Haematoxylin & Eosin staining and Oil Red O staining as previously published ^41,42^. No inflamed tissue areas were used in this current study. Additional, fresh-frozen and paraffin-embedded human samples were obtained from the Neuropathology Service at Brigham and Women’s Hospital and from archived autopsy specimens in the Department of Pathology and Laboratory Medicine of The Ottawa Hospital. Human spinal cord and muscle tissues were collected at The Ottawa Hospital with approval from the Ottawa Health Science Network Research Ethics Board.

Wild-type C57Bl/6J mice were used for analysis of the effects of *post mortem* interval on murine parkin in the brain. Mice from 4 to 8 months old were perfused with PBS and their brains were collected for PMI experiments.

### Sequential extraction of parkin from tissue

Roughly 1 cm^3^ samples of human brain frontal cortex and midbrain (age range 5-85 years of age) were weighed and placed in 3X volume/weight of Tris Salt buffer (TSS) (5mM Tris, 140 mM NaCl pH 7.5) containing complete EDTA-free protease inhibitor cocktail, and 10 mM iodoacetamide (IAA). The samples were homogenized on ice in a Dounce glass homogenizer by 50 passes, transferred to ultracentrifuge tubes and spun at 55,000 rpm and 4°C for 30 mins. The TS supernatant was transferred to a fresh tube and the pellet was extracted further with addition of 3x volume/weight of Triton X-100 buffer (TX, TS + 2 % Triton X-100). The samples were mixed by vortex, incubated on ice for 10 min and centrifuged again using the same prior setting. The TX supernatant was transferred to a fresh tube and the pellet was extracted further with addition of 3x volume/weight of SDS buffer (SDS, TS + 2 % SDS). The samples were mixed by vortex, incubated at room temperature for 10 min and centrifuged again at 55,000 rpm and 12°C for 30 mins. The SDS supernatant was transferred to a fresh tube and the pellet was either stored at −80°C or extracted further with addition of 3X volume/weight of 6X non-reducing Laemmli buffer (LB, 30 % SDS, 60 % glycerol, 0.3 % bromophenol blue, 0.375 M Tris. pH 6.8, 100mM DTT), mixed by vortex and incubated at room temperature for 10 min. Samples were centrifuged again at 55,000 rpm and 12°C for 30 mins and the LB supernatant was transferred to a fresh tube. Extracted proteins from TS, TXS and SDS buffers including pellet (20-30 µg) and 10-20 μL of LB extracts were run on SDS-PAGE using reducing (100 mM dithiothreitol, DTT) and/or non-reducing (0 mM DTT) loading buffer. Following transfer to membranes, Ponceaus S staining was used to confirm loading, and samples were blotted for parkin (Biolegend 808503, 1: 5,000), DJ-1 (ab18257, 1: 2,000), α-synuclein (syn1 1:1,000 or MJFR1 1:2000), LC3B (3868 1:2000), VDAC (MSA03 1:5000), MnSOD and GLO1 (1:1000), calnexin (MAB3126), cathepsin D (sc-6486), GRP75 (sc-1058). ImageJ software (1.52 k, National Institutes of Health, USA) was used for signal quantification purposes.

### mRNA Analysis

*PRKN* mRNA isolated from individual *S. nigra* DA neurons (SNDA), cortical pyramidal neurons (PY) and non-neuronal, blood mononuclear cells (NN) were processed, as described ^43^ and as annotated in the Human BRAINcode database (www.humanbraincode.org).

### ROS (H_2_O_2_) measurements

Amplex^®^ red hydrogen peroxide/peroxidase assay kit (Invitrogen A22188) was used to monitor endogenous levels of H_2_O_2_. Pre-weighed cortex pieces from human brains were homogenized on ice in the 1x reaction buffer provided, using a Dounce homogenizer (3 times volume to weight ratio). Homogenates were diluted in the same 1x reaction buffer (10x and 5x). A serial dilution of the H_2_O_2_ standard provided was prepared (20, 10, 2 and 0 µM). 50 µL of standards and samples were plated in a 96 well black plate with clear flat bottom. The reaction was started by the addition of 50µL working solution which consist of 1x reaction buffer, Amplex^®^ red and horseradish peroxidase. The plate was incubated at room temperature for 30 minutes protected from light. A microplate reader was used to measure either fluorescence with excitation at 560 nm and emission at 590 nm, or absorbance at 560 nm. The obtained H_2_O_2_ levels (µM) were normalized to the tissue weight (g).

### Recombinant protein expression in pET-SUMO vector

Wild-type and truncated (residues 321-465) human parkin were expressed as 6His-Smt3 fusion proteins in *Escherichia coli* BL21 (DE3) Codon-Plus RIL competent cells (C2527, New England Biolabs) as previous described ^44-46^. DJ-1 and SNCA coding sequences were cloned from a pcDNA3.1 vector into the pET-SUMO vector using PCR and restriction enzymes. ARPD-associated parkin mutants in the pET-SUMO vector were generated using site-directed mutagenesis. All proteins were overexpressed in Escherichia coli BL21 Codon-Plus competent cells (C2527, New England Biolabs) and grown at 37 °C in 2 % Luria Broth containing 30 mg/L kanamycin until OD600 reached 0.6, at which point the temperature was reduced to 16°C. All Parkin-expressing cultures were also supplemented with 0.5 mM ZnCl_2_. Once OD600 reached 0.8, protein expression was induced with isopropyl β-D-1-thiogalactopyranoside, except ulp1 protease, which was induced once OD600 had reached 1.2. The concentration of IPTG used for each construct is as follows: 25 µM for wild-type and point mutants of parkin, and 0.75 mM for truncated parkin, DJ-1, α-synuclein, SAG, and ulp1 protease. Cultures were left to express protein for 16-20 h. Cells were then harvested, centrifuged, lysed and collected on Ni-NTA agarose beads in elution columns.

### Protein redox chemistry and oxidative toxin conjugation

The recombinant protein samples were first prepared by removing excess TCEP, present in the elution buffer using repeat centrifugations (8 times 4000 x g at 4°C for 10 min) in Amicon Ultra 10kDa MWCO filters. The protein concentrations were measured and adjusted to 20µM. Stock solutions of rotenone (1 mM), carbonyl cyanide m-chlorophenyl hydrazone (CCCP, 50 mM), and hydrogen peroxide (H_2_O_2_, 9.8 mM) were prepared. Aminochrome was freshly synthesized from Dopamine (see below). An aliquot of 10µL of each protein sample (at 20 µM) was reacted with oxidants at the following concentrations: 0, 10, 20, 200, 2000 µM rotenone; 0, 10, 20, 200 µM CCCP; 0, 2, 20, 50, 200 aminochrome; 0, 20, 200, 500, 750, 1000, 2000 µM H_2_O_2_ or 0, 10, 50, 100, 200, 500, 1,000 μM DTT. The samples were treated for 30 min at 37°C and centrifuged at 14,000 rpm for 15 min. The supernatant was transferred to a fresh tube and the remaining pellet was extracted with 10µL of T200-TCEP containing either 10 % SDS or 100 mM DTT. The pellets were incubated again for 30 min at 37°C and centrifuged at 14,000 rpm for 15 min. Laemmli buffer (10 µL, containing 100 mM mercaptoethanol) was added to both the pellet and supernatant fractions and samples were separated on two SDS-PAGE. One gel was used for in-gel protein staining and the other was used for NBT staining. Specific bands of aminochrome treated wild-type full length r-parkin were excised from silver-stained gels and analyzed by LC-MS/MS as described below.

### Aminochrome synthesis

A solution of 0.1 M sodium phosphate buffer pH 6.0 was prepared from a mixture of 12 mL of 1M NaH_2_PO_4_ and 88.0 mL of 1M Na_2_HPO_4_. The reaction buffer (0.067 M sodium phosphate, pH 6.0) was prepared by adding 33 mL of 0.1 M sodium phosphate buffer to 17 mL water. A solution of 10mM dopamine in reaction buffer was prepared by adding 19 mg of dopamine hydrochloride to 1 mL of reaction buffer. Oxidation was activated by adding 5 µL of tyrosinase (25,000 U/mL) and the mixture was incubated at room temperature for 5 min. The tyrosinase was separated from the oxidized dopamine using a 50 kDa cut-off Amicon Ultra centrifugation filter by centrifuging at 14,000 rpm for 10 min. The absorbance of the filtrate was measured at a wavelength of 475 nm using Ultrospec 21000 pro spectrophotometer and the concentration of aminochrome was determined using the Beer-Lambert equation and extinction coefficient of 3058 L x mol^-1^ × cm^-1^.

### Protein staining methods

All proteins were separated on pre-cast 4-12 % Bis-Tris SDS-PAGE gels (NPO321BOX, NPO322BOX, NPO336BOX) from Invitrogen using MES running buffer (50mM MES, 50mM Tris, 1mM EDTA and 0.1 % SDS, pH 7.3) and Laemmli loading buffer (10% SDS, 20% glycerol, 0.1% bromophenol blue, 0.125M Tris HCl, 200mM DTT or β-mercaptoethanol). Proteins were stained in gel using SilverQuest™ Silver Staining Kit (LC6070) from Invitrogen or Coomassie brilliant blue R-250 dye (20278) from ThermoFisher Scientific using the following protocol: The gel was transferred to a plastic container and rocked for 30 min in Fix Solution (10% acetic acid, 50% methanol), followed by staining for 2-24 h (0.25% Coomassie R250) until the gel turned a uniform blue. The stain was replaced with Destain Solution (7.5% acetic acid and 5% methanol) and the gel was rocked until crisp blue bands appeared. Following a wash with water the gel was stored in 7 % acetic acid. Proteins transferred to PVDF (1620177, Bio-Rad) membranes were stained with Ponceau S solution for 20 min, washed three times with water, imaged and then destained with 0.1M NaOH prior to Western blotting. Proteins conjugated to redox-cycling molecules were stained using the following NBT protocol: Following Ponceau S staining and NaOH destaining, the membrane was washed twice with water, covered from light and incubated with rocking for 45 min in NBT solution [8-9 mg of NBT added to 14 mL of 2 M potassium glycinate buffer (75 g glycine in 0.4 L water and pH adjusted to 10.0 with KOH)]. The membrane was washed twice with destain solution (0.16M sodium borate) and incubated in fresh destain solution for 16-24 h. The membrane was washed once with water and imaged before drying.

### Dynamic light scattering (DLS)

The protein buffer (50mM Tris, 200mM NaCl and 250µM TCEP, pH 7.5) was exchanged for a 20mM phosphate buffer with 10mM NaCl (pH 7.4). 20 µM full-length wild-type r-parkin was centrifuged at 14,000 rpm for 60 min at 4 °C and light scattering intensity of the supernatant was collected 30 times at an angle of 90° using a 10 sec acquisition time. Measurements were taken at 37 °C using a Malvern Zetasizer Nano ZS instrument equipped with a thermostat cell. The correlation data was exported and analyzed using the nanoDTS software (Malvern Instruments). The samples were measured at 0-, 1-, 3- and 5 hours. Following 24 hr incubation, 2 mM DTT was added to the sample and the light scattering intensity of the supernatant was measured again.

### Far UV circular dichroism spectroscopy

15 µM of reduced and partially oxidized full-length wild-type r-parkin was measured at t = 0 and t = 5 days of incubation under native conditions in 20 mM phosphate, 10 mM NaCl buffer. The aggregates rich phase and the monomer rich phase in the samples were separated with ultracentrifugation (100,000 g for 2 hours). Far UV circular dichroism (CD) spectra were recorded for the monomer and aggregated rich phase of protein samples using a JASCO J-720 spectrometer. The final spectrum was taken as a background-corrected average of 5 scans carried out under the following conditions: wavelength range 250–190 nm at 25 °C; bandwidth was 1 nm; acquisition time was 1 sec and intervals was 0.2 nm. Measurements were performed in a 0.01 cm cell. CD spectra were plotted in mean residue molar ellipticity units (deg cm^2^ dmol^-1^) calculated by the following equation: [Θ] = Θ_obs_/(10*ncl)*, where [Θ] is the mean residue molar ellipticity as a function of wavelength, Θ_obs_ is the measured ellipticity as a function of wavelength (nm), *n* is the number of residues in the protein, *c* is the concentration of the protein (M), and *l* is the optical path length (cm). Secondary structure analysis of proteins using CD spectroscopic data was carried out using the BeStSel (Beta Structure Selection) software ^47,48^.

### Chemiluminescence/ direct reactive oxygen species (ROS) assay

The assay was modified from Muller et al. 2013 to measure the ROS-quenching ability of Parkin proteins, DJ-1, SNCA, BSA, GSH, and catalase. Protein concentrations were quantified using Bradford assay and adjusted to 5, 10, 15 and 30 µM in buffer not containing TCEP. BSA (10 and 20 µM), GSH (15, 20, 200, 400, 800 and 2000 µM), and catalase (0.015, 0.15, 0.25 and 15 µM) were prepared. Stock solutions of H_2_O_2_ for standard curve were prepared at 5, 10, 20, 40 and 50 mM in 0.1 M Tris HCl pH 8.0 using 30 % H_2_O_2_. Stock solutions of 300 mM luminol and 40 mM 4-iodophenol were prepared in DMSO and protected from light. Signal reagent, containing 1.94 mM luminol and 0.026 mM 4-iodophenol, was prepared in 0.1 M Tris HCl pH 8.0 and protected from light. A 0.4 % horseradish peroxidase solution was prepared using HRP-linked anti-rabbit secondary antibody diluted in Stabilizyme solution (SurModics SZ02). Each read was set up in triplicate on a white polystyrene 96-well plate (ThermoFisher 236105) and to each well was added 80 µL Stabilizyme, 15 µL of 0.4 % horseradish peroxidase (HRP) and 25 µL of sample or controls. One of the injectors in a Synergy H1Multi-Mode Plate Reader (Bio Tek) was primed and set to inject 15 µL of signal reagent and 15 µL of each H_2_O_2_ stock solution was manually added to corresponding controls and samples just prior to reading. Final concentrations of reagents were 0.04 % HRP, 500, 1000, 2000, 4000 and 5000 µM H_2_O_2_, 194 µM luminol, 2.6 µM 4-iodophenol and 0.8, 1.7, 2.5 or 5 µM of protein. The plate reader was set to measure luminescence every 1 min for a total of 10 min. The resulting kinetic data was converted to area under the curve (AUC) using Prism version 6. For samples pre-incubated with 20 mM iodoacetamide, a stock solution of 1 M iodoacetamide was prepared. To each well containing 25 µL of sample, 0.52 µL of 1 M iodoacetamide and 0.48 µL of buffer not containing TCEP was added and the samples were incubated for 2 h at 37°C. Following incubation, the reagents for chemiluminescence were added as above except 79 µL of Stabilizyme was used instead of 80 µL and the samples were analyzed as above.

### Thiol quantification of recombinant proteins

Recombinant protein samples were first prepared by exchanging the T200 protein buffer (50 mM Tris, 200 mM NaCl and 250 µM TCEP, pH 7.5) for T200-TCEP using repeat centrifugations (8 times 4000 x g at 4°C for 10 min) in Amicon Ultra 10 kDa MWCO filters. The protein concentrations were measured and recorded. The glutathione stock solution of 32,539 µM was prepared by dissolving 1 mg glutathione (GSH) in 1 mL of T200-TCEP and the standards 0, 50, 101, 203, 406, 813 and 1000 µM were prepared by serial dilution in T200-TCEP. The reaction buffer (0.1 M sodium phosphate, pH 8.0) was prepared by adding 93.2 mL 1M Na_2_HPO_4_ and 6.8 mL of NaH_2_PO_4_ in 1 L of water. Thiol detecting reagent (Ellman’s reagent) was prepared by dissolving 2 mg of 5,5’-dithio-bis-[2-nitrobenzoic acid] (DNTB) in 1 mL of reaction buffer. The assay was performed in 96-well clear round bottom plates by adding 50 µL of thiol detecting reagent to 50 µL of sample or standard and incubating for 15 min at room temperature. The resulting 5-thio-2-nitrobenzoic-acid (TNB) produced was measured by absorbance at 412 nm using a Synergy H1Multi-Mode Plate Reader (Bio Tek). The amount of free thiols detected in each sample was calculated using the regression curve obtained from the glutathione standards and dividing by the concentration of the sample.

### Cysteine labeling for mass spectrometry

The recombinant protein samples were first prepared by exchanging the T200 buffer for PBS. The protein concentrations were measured and adjusted to 10 µM using PBS. Stock solutions of 500 mM DTT, 100 mM iodoacetamide (IAA), 100 mM hydrogen peroxide and 250 mM ethylenediaminetetraacetic acid (EDTA) were prepared in PBS. A stock of 500 mM N-ethyl-maleimide (NEM) was prepared in ethanol immediately before use. For the first optimization and comparison of IAA and NEM labelling (*i.e.*, Table S2), r-parkin was treated with 2 mM DTT for 30 min at 37°C followed by incubation with 5 mM IAA or 85 mM NEM for 2 h at 37°C. The stepwise cysteine labeling procedure was as follows: A 10 µL aliquot of protein (at 10 µM) was reacted with hydrogen peroxide at various concentrations, as indicated (Table 1) for 30 min (and up to 60 min) at 37°C as indicated. Any unreacted cysteines were alkylated with incubation with 5 mM IAA (either with or, in some runs, without 10 mM EDTA) for 2 hrs at 37°C. Previously oxidized cysteines were then reduced by treatment with 40 mM DTT for 30 min at 37°C. Newly reduced cysteines were alkylated by incubation with 85 mM N-ethyl maleimide (NEM) for 2 hrs at 37°C. The samples were separated on SDS-PAGE using Laemmli buffer containing 100 mM DTT and proteins visualized using Coomassie staining. Appropriate bands were excised and analyzed by liquid chromatography mass spectrometry (LC-MS/MS).

### Protein identification by LC-MS/MS

Proteomics analysis was performed at the Ottawa Hospital Research Institute Proteomics Core Facility (Ottawa, Canada). Proteins were digested in-gel using trypsin (Promega) according to the method of Shevchenko^49^. Peptide extracts were concentrated by Vacufuge (Eppendorf). LC-MS/MS was performed using a Dionex Ultimate 3000 RLSC nano HPLC (Thermo Scientific) and Orbitrap Fusion Lumos mass spectrometer (Thermo Scientific). MASCOT software version 2.6.2 (Matrix Science, UK) was used to infer peptide and protein identities from the mass spectra. For detection of dopamine metabolites on Parkin, the following variable modifications were included: 5,6-indolequinone (+C_8_O_2_NH_3_, m/z shift +145), aminochrome (+C_8_O_2_NH_5_, +147), aminochrome +2H (+C_8_O_2_NH_7_, +149), and dopamine quinone (+C_8_O_2_NH_9_, +151). These samples were prepared for analysis without any use of dithiothreitol or iodoacetamide. The observed spectra were matched against human sequences from SwissProt (version 2018-05) and also against an in-house database of common contaminants. The results were exported to Scaffold (Proteome Software, USA) for further validation and viewing. Analysis of the holoprotein and of three runs of H_2_O_2_-exposed r-parkin (Supplemental Table 2) were performed at the University of Western Ontario. There, samples were run on a QToF Ultima mass spectrometer (Waters) equipped with a Z-spray source and run in positive ion mode with an Agilent 1100 HPLC used for LC gradient delivery (University of Western Proteomics Facility).

### MaxQuant analysis of mass spectrometry data

For applicable experiments, the raw MS data files were further processed with MaxQuant software version 1.6.5 and searched with the Andromeda search engine^18^. The reference fastas were set to uniprot-human (version 2019-02-12) and uniport-ecoli. The E. coli proteome was included to account for bacterial proteins present in the recombinant protein samples. The ‘second peptides’ and ‘match between runs’ settings were enabled. All other settings were left as default. Selected variable modifications included oxidation (Met), acetylation (protein N-terminus), and carbamidomethyl (Cys), as well as custom modifications for pyro-carbamidomethyl (N-terminal Cys), N-ethylmaleimide (Cys), and NEM+water (Cys). For data analysis, site-level intensity values were obtained from the MaxQuant-generated “CarbamidomethylSites” table which combines the intensity of MS1 signals from all peptides covering a particular cysteine residue.

### Immunoprecipitation (IP) of brain parkin

Conjugation of anti-parkin antibody (Prk8, 808503, lot B209868) to magnetic beads at a final concentration of 10 mg of antibody/ mL of beads was done following the Magnetic Dynabeads Antibody Coupling Kit from Invitrogen (14311D). Human tissue lysates were also prepared using the “Sequential Extraction of Proteins from Tissue” protocol as described above with addition of 10 mM iodoacetamide prior to homogenization. TS tissue extracts (n=4) and SDS tissue extracts (n=8) were diluted in TS buffer, resulting in a final SDS concentration of 0.0175 % and 0.05 % respectively. For the IP, Prk8 conjugated agarose beads were first prepared by multiple washes with 1 mL of TS buffer using centrifugation (1000 x g at 4°C for 3 min) and adhesion to a strong magnet. Amounts of Prk8 conjugated agarose beads used for each experiment were approximated based on the amount of parkin (µg) / sample calculated by densitometry when the sample was compared to recombinant parkin protein standards using Western blotting with Prk8 primary antibody. The mixture was incubated for 16 h at 4°C with slow rotation. Unbound proteins, which did not bind to the Prk8 conjugated agarose beads, were separated from the beads by centrifugation (1000 x g at 4°C for 3 min) followed by adhesion to a strong magnet and saved as the IP “unbound” fraction. Beads from cellular or human IP were washed three times with 900 or 1000 µL respectively of ice-cold RIPA buffer (1 % nonionic polyoxyethylene-40, 0.1 % SDS, 50 mM Tris, 150 mM NaCl, 0.5 % sodium deoxycholate, 1 mM EDTA) using centrifugation (1000 x g at 4°C for 3 min) and adhesion to a strong magnet. Approximately 5-10 µL of each wash was combined and saved as the IP “wash” fraction. To elute Prk8 bound proteins, 15-35 uL of 6X reducing Laemmli buffer (30 % SDS, 60 % glycerol, 0.3 % bromophenol blue, 0.375 M Tris, 100 mM DTT, pH 6.8) was added to the beads and the samples were boiled for 5 min. Following centrifugation (1000 x g at 4°C for 3 min), the supernatant was transferred to a fresh tube labeled “IP elute” and the beads were discarded. To assess IP efficiency, eluted fractions (IP elute), along with controls (input, unbound, wash and recombinant parkin protein standards) were run on SDS/PAGE and blotted with anti-parkin (MAB5512 or 2132S). Human IP elutes used in subsequent for mass spectrometry (MS) analysis were incubated with 500 mM N-ethyl maleimide (as indicated for select runs) for 16 h at 4°C prior to SDS-PAGE and further processing for MS (as described above). Gel slices corresponding to band sizes 50-75 kDa were excised and analyzed by LC-MS/MS.

### *In vitro* melanin formation assay

The recombinant protein samples were first prepared by exchanging the T200 protein buffer (50 mM Tris, 200 mM NaCl and 250 µM TCEP, pH 7.5) for T200-TCEP (50 mM Tris and 200 mM NaCl, pH 7.5) using repeat centrifugations (8 times 4000 x g at 4°C for 10 min) in Amicon Ultra 10 kDa MWCO filters. The protein concentrations were measured and adjusted to 20 µM using T200-TCEP. A 0.067 M sodium phosphate buffer, pH 6.0, was prepared by adding 33 mL of 0.1 M sodium phosphate buffer to 17 mL water and adjusting the pH using HCl. A stock solution of 100 mM dopamine HCl was prepared in 0.067 M sodium phosphate buffer and stock solutions of 100 mM reduced glutathione (GSH) and hydrogen peroxide were prepared in T200-TCEP.

Samples and controls were prepared in 100 µL total volume and contained: 10 µL of 20 µM protein or T200-TCEP, 10 µL of 100 mM dopamine or 0.067 M sodium phosphate buffer, 10 µL of 100 mM glutathione or T200-TCEP buffer, and 70 µL T200-TCEP. The final concentration of protein was 2 µM and the final concentration of reagents was all 10 mM. The samples and controls were plated in triplicate, and absorbance read at 405 and 475 nm every 90 sec for 1 h. The plate was then protected from light and incubated for 24 h. A final reading of absorbance at 405 and 475 nm was taken after 24 h.

### Cell cytotoxicity assay

Human neuroblastoma cell line (M17 cells) wild-type, Vector (Myc), P5 (low stable expression of Myc-parkin) and P17 (high stable expression of Myc-parkin) were grown in 6 well culture plates, at 0.3×10^6^ cell density (80% confluence) in Opti-MEM media (Gibco 11052-021) containing heat inactivated FBS (Gibco 10082-147), Pen/strep/Neo (5mg/5mg/10mg) (Gibco 15640-055), MEM non-essential amino acids (10mM) (Gibco 11140-050) and sodium pyruvate (100mM). For rescue experiments, flag-vector, flag-parkin, flag-p.G328E and flag-p.C431F pcDNA3.1+ were overexpressed in M17 wild-type cells. 4 µg of cDNA was transfected using a 1:1 ratio of cDNA: Lipofectamine 2000 (52887, Invitrogen) in OPTI-MEM transfection medium. The cDNA and Lipofectamine 2000 was first incubated for 20 min at room temperature before being applied directly to the cells for 1 h at 37°C with 5 % CO_2_ followed by direct addition of fresh growth medium. The cells were incubated another 24 hours at 37°C with 5 % CO_2_.

Dopamine hydrochloride (Sigma H8502) 200 mM stock was prepared. The cells were washed with fresh media once and then incubated with media alone or supplemented with dopamine at final concentrations of 20 µM and 200 µM for 18-20 hours. Post dopamine stress, media was collected from all wells for cytotoxicity assay, the cells were harvested and lysed with TS buffer and centrifuged. The supernatant was collected and saved for western blot analysis and to assess total cell toxicity signal. The pellet was suspended in SDS buffer and centrifuged.

Vybrant ™ cytotoxicity assay kit (Molecular Probes V-23111) was used to monitor cell death through the release of the cytosolic enzyme glucose 6-phosphate dehydrogenase (G6yPD) from damaged cells into the surrounding medium. 50 µl of media alone (no cells), media from control and stressed cells and cell lysates were added to a 96-well microplate. Fifty µl of reaction mixture, containing reaction buffer, reaction mixture and resazurin, was added to all wells, and the mircroplate was incubated at 37°C for 30 mins. A microplate reader was used to measure either fluorescence with excitation at 560 nm and emission at 590 nm. A rise in fluoresence indicates a rise in G6PD levels i.e. a rise in cell death.

### Immunohistochemistry (IHC)

Immunohistochemistry was performed on paraffin-embedded sections and treated as previously described (Schlossmacher et al. 2002; Schlossmacher and Shimura 2005, Shutinoski et al 2019). Briefly, prior to antibody incubation, sections were deparaffinized in xylene and successively rehydrated through a series of decreasing ethanol concentration solutions. Endogenous peroxidase activity was quenched with 3% hydrogen peroxide in methanol, followed by a standard citric acid-based antigen retrieval protocol to unmask epitopes. Sections were blocked in 10-20% serum in PBS-T to reduce non-specific signal. Sections were incubated overnight at 4°C in primary antibodies diluted in 1-5% serum in PBS-T according to the following concentrations: novel Parkin mAbs from Biolegend clones D (BioLegend, A15165D; 1:250), clone E (BioLegend, A15165E; 1:2000), and clone G (1:250), PRK8 (BioLegend, MAB5512; 1:500), Lamp3 (Santa Cruz, SC5275; 1:100), LC3B (Sigma, L7543-200uL; 1:100), VDAC (MitoScience, MSA03; 1:100). Biotinylated secondary antibodies (biotinylated anti mouse IgG (H+L) made in goat; Vector Labs, BA-9200, biotinylated anti-rabbit IgG (H+L) made in goat; Vector Labs, BA-1000) were diluted to 1:225 and sections were incubated for 2 hours at room temperature. The signal was amplified with VECTASTAIN® Elite® ABC HRP Kit (Vector Labs, PK-6100), and visualized via standard DAB solution, 55mM DAB, or Vina green (Biocare Medical, BRR807AH), or most commonly metal enhanced DAB (Sigma, SIGMAFAST™ DAB with Metal Enhancer D0426). Samples were counterstained with Harris Modified Hematoxylin nuclei stain and dehydrated through a series of increasing ethanol concentration solutions and xylene. Permount (Fisher Scientific, SP15-100) was used for mounting and slides were visualized with high magnification images via a Quorum Slide Scanner (Ottawa Hospital Research Institute).

### Immunofluorescence (IF) and confocal microscopy

Paraffin-embedded human midbrain sections were stained by routine indirect immunofluorescence with the following details. Antigen retrieval was performed in Tris-EDTA buffer pH 9 for 10 mins. Primary antibodies were incubated overnight at 4C. Details for primary antibodies anti-Parkin Clone E (1:500), anti-LAMP3 (1:250) are described above. A 40 minute incubation with the following secondary antibodies was performed: goat anti-mouse alexa fluor 488 (1:200), goat anti-rabbit alexa fluor 594 (1:500). Slides were mounted with fluorescence mounting medium with DAPI. Stained sections were imaged using a Zeiss LSM 880 AxioObserver Z1 with an Airyscan Confocal Microscope then processed and analyzed using Zeiss Zen and Fiji software.

### Statistical analyses

All statistical analyses were performed using GraphPad Prism version 6 (GraphPad Software, San Diego, CA, USA, www.graphpad.com). Differences between two groups were assessed using an unpaired t-test. Differences among 3 or more groups were assessed using a one-way or two-way ANOVA followed by Tukey’s post hoc corrections to identify statistical significance. Subsequent post hoc tests are depicted graphically and show significance between treatments. For all statistical analysis a cut-off for significance was set at 0.05. Data is displayed with p values represented as *p < 0.05, **p < 0.01,***p < 0.001, and ****p < 0.0001.

#### Logistic/linear regression modelling

Linear regression (for continuous dependent variable, e.g. H_2_O_2_ level, mRNA level) or logistic regression (for binary dependent variable, *e.g.*, parkin present in TS fraction) modelling were performed. Furthermore, to address the effect of age on parkin solubility, receiver operating characteristic (ROC) curve and area under the ROC curve (AUC) were calculated, as reported^50^.

## Supporting information

Tokarew et al_Supplementary Tables

## ACKNOWLEDGMENTS

We are grateful for the commitment of patients and their family members to participate in autopsy studies. We thank Dr. J. Palacino for creating stable M17 cell lines, Drs. A. Brice and E. Fon for sharing *prkn*-null mice, Dr. B. Madras for providing specimens of cynomolgus brain, Drs. R. Tam, L. Dong and Ms. K. Solti and Ms. H. Boston for technical support, Dr. D. Gibbings for antibodies, Dr. D. Gray for assistance with confocal imaging, Drs. M. Medina and R. R. Ratan for encouragement, Drs. S. Bennett, D. Pratt for discussions, and Drs. L. Zecca, F. Zucca, H. Lochmueller, M. Rousseaux, and members of the Schlossmacher lab for their critical comments.

## Author contributions

*Study design:* J.M.T., D.N.E.K., P.T., J.J.T., M.G.S.; *Writing and Figure preparation:* J.M.T., D.N.E.K., N.A.L., T.K.F., M.J., A.P.N., J.L., G.S.S., J.M.W., G.T., P.T., J.J.T., and M.G.S. prepared the initial draft of the manuscript and figures. All authors reviewed and / or edited the manuscript and approved of the submitted versions. *Experiments*: J.M.T., D.N.E.K., N.A.L., T.K.F., M.J., A.P.N., B.O., L.W., J.K., A.C.N., Q.J., R.S., J.L., M.Z., K.R.B., A.T., X.D., L.P., G.T. performed experiments; and C.R.S., A.B.W., E.T., A.H., A.P., J.A.C., provided data, tissue specimens and critical comments. *Analysis:* J.M.T., D.N.E.K., J.L., T.K.F., G.S.S., L.P., G.T., J.M.W., P.T., J.J.T., M.G.S. performed data analyses. *Study supervision*: P.T., J.J.T., M.G.S. *Overall responsibility:* M.G.S. This work is dedicated to the memory of Mr. Bruce Hayter (1962-2019), a tireless advocate for persons with young-onset parkinsonism.

## Data and materials availability

Original data associated with this study are available in the main text and extended data figures and tables; additional data will be made available upon request.

---

**Extended Data Figure 1.**
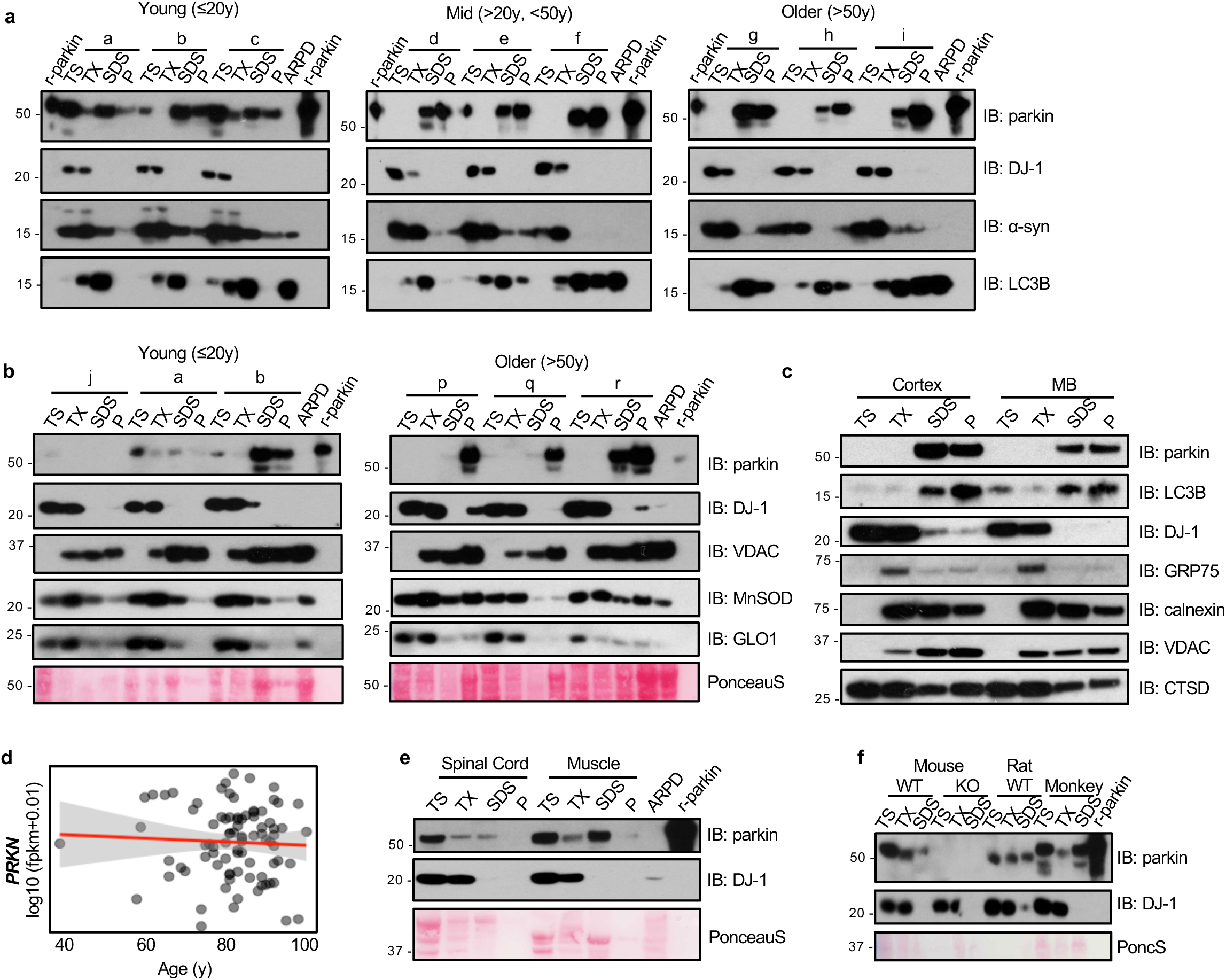
Parkin transitions from a soluble to an aggregated state in adult human brain. **(a)** Western blots of parkin, DJ-1, α-synuclein and LC3B distribution in 9 representative human cortex samples identified by lowercase letter code (see **Supplementary Information Table 1**). Tissue fractionation and age ranges were as described in Fig. 1; gels were run under reducing conditions. ARPD lysates and r-parkin are included as controls. **(b)** Western blots of parkin, DJ-1, VDAC, MnSOD and glyoxylase-1 proteins, and Ponceau S staining in serially fractionated human cortices from younger (n=3) and older (n=3) individuals. Quantification of relative protein distribution is shown in **Fig. 1h**. **(C)** Western blot of indicated proteins from serially fractionated cortex and midbrain tissue as described in (a). **(d)** Linear regression analysis of log transformed *PRKN* transcripts as a function of age in human control *S. nigra* dopamine neurons where each dot represents values for a single neuron as shown in **Fig. 1e**. **(e)** Western blot of parkin and DJ-1 expression in serial fractions from representative human spinal cord and skeletal muscle tissues from individuals ≥ 50y. Relative protein distribution is shown in **Fig. 1i**. **(f)** Western blot of parkin and DJ-1 and Ponceau S staining of serial fractions from whole brains of wild-type (WT; 8 mths of age) and *Prkn* knock-out (KO) mice, rat (WT; 14 mths) and cortex from cynomolgus monkey (60 mths).

**Extended Data Figure 2.**
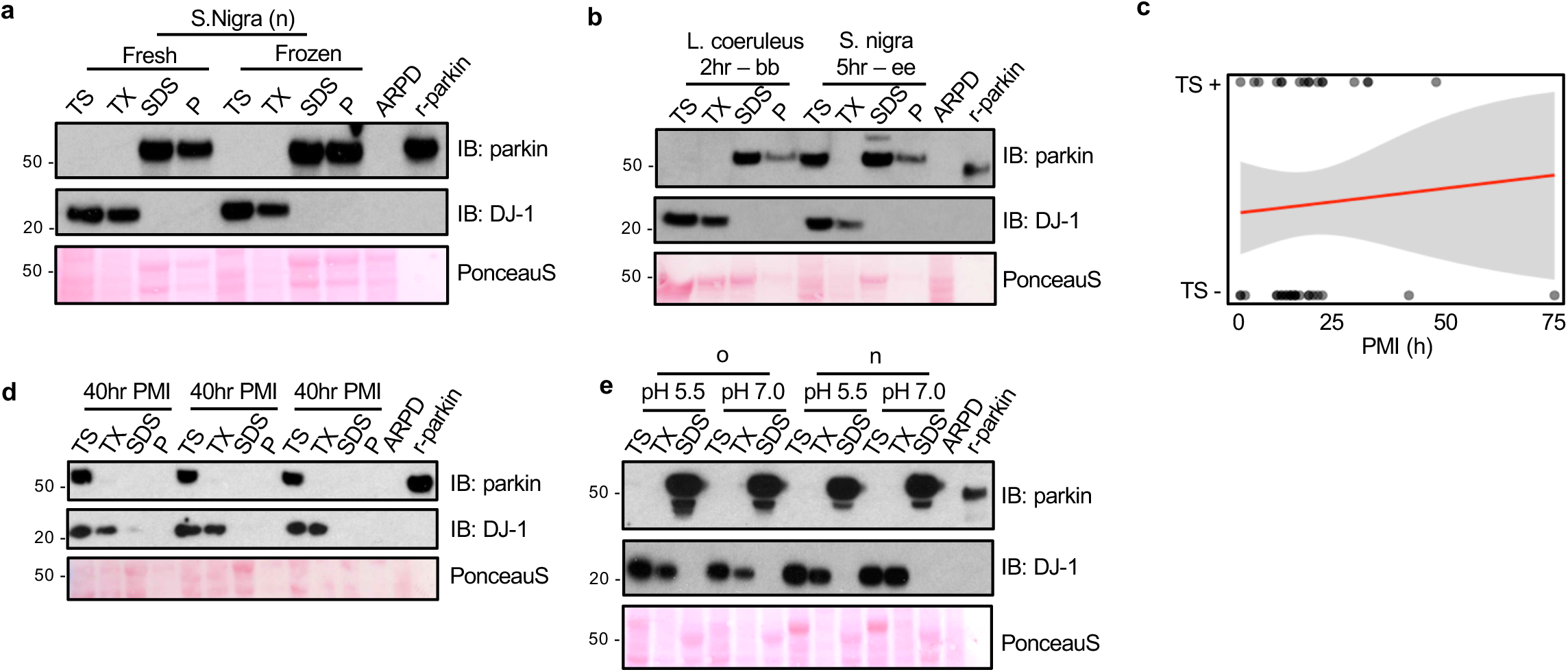
Parkin solubility is not affected by tissue freezing, length of *post mortem* interval or pH changes. **(a-b)** Western blot of parkin and DJ-1 distribution of **(a)** human *S. nigra* (brain code n) lysates that had been processed in parallel with *vs.* without a freeze/thaw step prior to serial fractionation, and **(b)** from *L. coeruleus* and *S. nigra* which were collected within 2-5 hrs after death prior to freezing. **(c)** Logistic regression analysis of parkin solubility as a function of length for *post mortem* interval (PMI; in hrs); the logistic regression analysis line (red) and 95% confidence intervals (grey) are shown (n=45 cortices). **(d)** Immunoblots for endogenous parkin and dj-1 signals from serially extracted, WT mouse brains (n=3) dissected following 40 hr *post mortem* interval at 4°C. **(e)** Western blotting of parkin and DJ-1 distribution and Ponceau S staining for fractions of human cortices serially extracted in parallel using standard buffers with varying pH, as indicated.

**Extended Data Figure 3.**
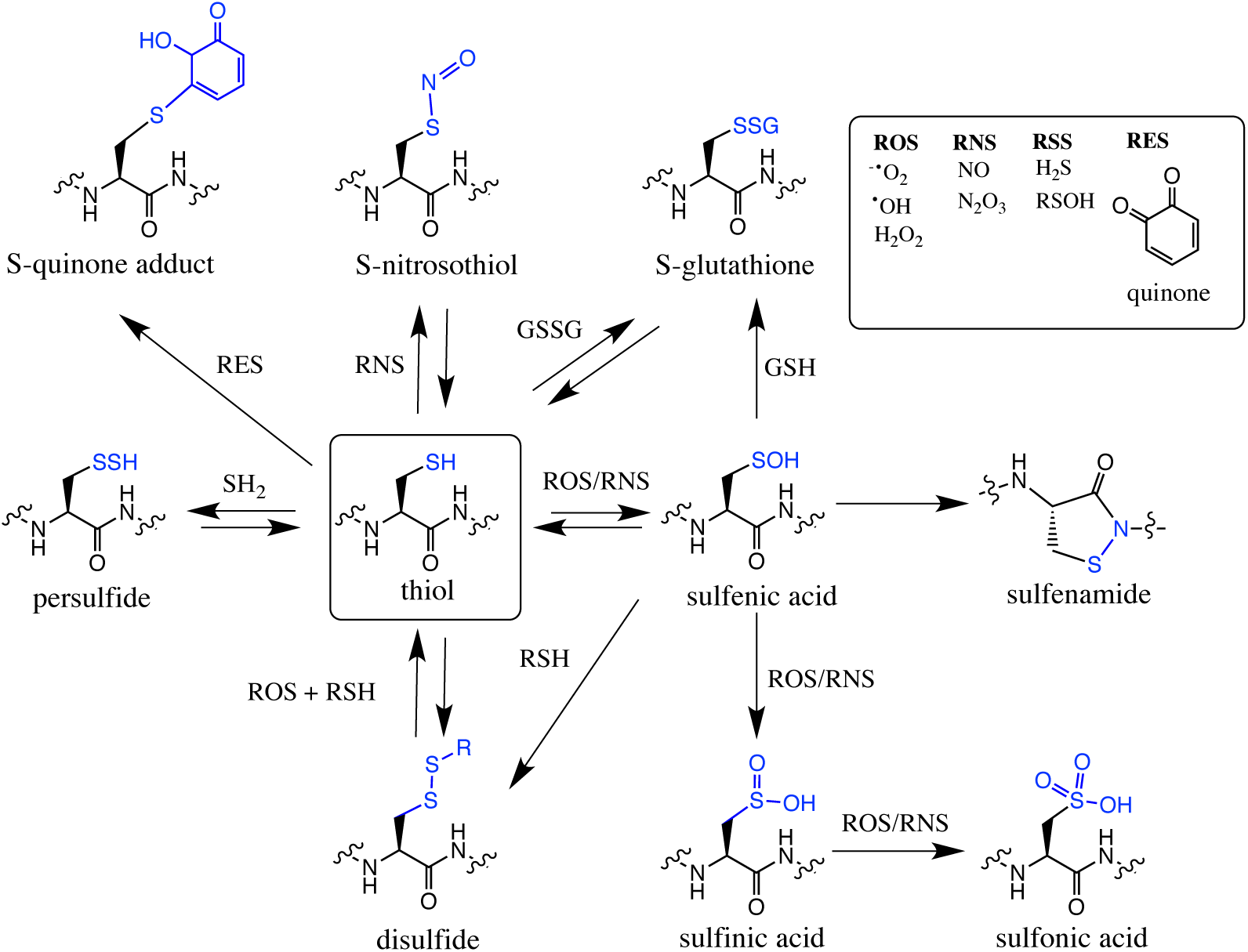
Summary of redox-related thiol chemistry. Schema of select, reversible and irreversible cysteine modifications that can occur on thiols due to attacks by reactive oxygen species (ROS), -nitrogen species (RNS), -sulfur species (RSS) and -electrophilic species (RES), which include quinones. This graphic summary was modified from Alcock *et al.*, 2018.

**Extended Data Figure 4.**
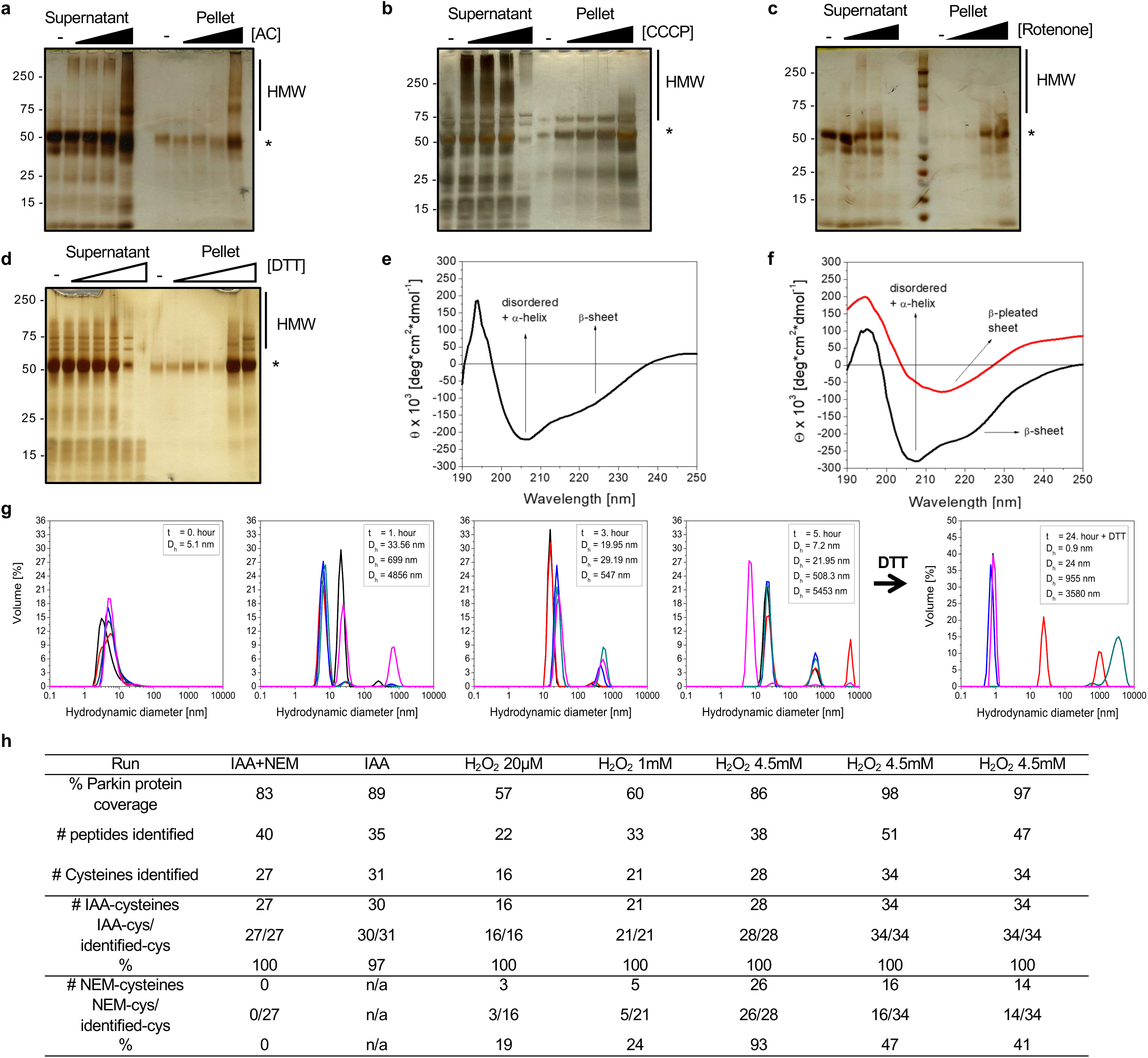
Oxidation of parkin thiols promotes insolubility. **(a-d)** Detection of r-parkin in soluble (supernatant) and insoluble phases (pellet; recovered by 10% SDS-containing buffer) of following exposure to increasing concentrations of **(a)** aminochrome (0-200 µM), **(b)** CCCP (0-200 µM), (**c**) rotenone (0-2mM) and **(d)** DTT (0-1M) as described in **Fig. 2a**. **(e-f)** Circular dichroism spectra of soluble, H_2_O_2_-exposed WT r-parkin secondary structure at **(e)** T=0 and **(f)** soluble (black line) and aggregated (red line) states following incubation at 37o C for T=5 days to promote spontaneous oxidation as described in **Fig. 2c,d**. **(g)** Dynamic light scattering analysis showing progressive size changes, as measured in hydrodynamic diameters (nm), was monitored during 5 hrs at room temperature. The structural state for wild-type, r-parkin under non-reducing, native conditions showed increased aggregate formation over time, which was partially reversed by treatment with DTT as indicated. **(h)** Summary of LC-MS/MS based analysis of cysteine oxidation state of untreated and H_2_O_2_-treated r-parkin as indicated using IAA-DTT-NEM fingerprinting to identify reduced cysteines (IAA) or reversibly-oxidized residues (NEM). For comparison, see **Extended Data Table 1** that lists modified cysteines identified in parkin purified from human brain cortex. A complete list of modified cysteines under each condition tested is found in **Supplemental Information Table 2**.

**Extended Data Figure 5.**
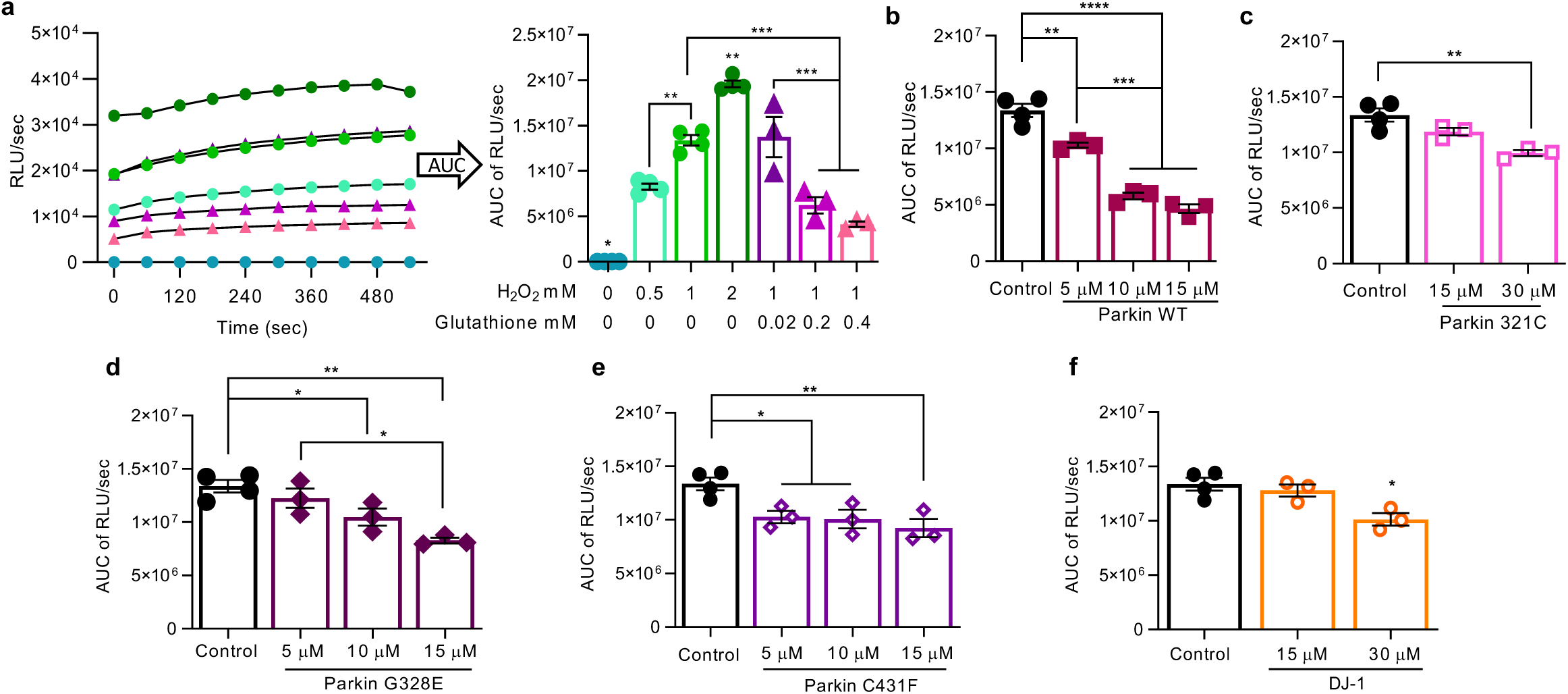
Parkin directly reduces H_2_O_2_ to water in a concentration-dependent manner. **(a)** Kinetic readings from *in vitro* colorimetric H_2_O_2_ assays comparing increasing concentrations of input H2O2 and the effect of increasing concentrations of glutathione. Curves were then converted to the area under the curve (AUC) where AUC integrates the total value of H_2_O_2_ signals generated over the 10 minutes time course of the assay. **(b-f)** AUC graphs for results from *in vitro* H_2_O_2_ assays for various concentrations of recombinant proteins, as indicated.

**Extended Data Figure 6.**
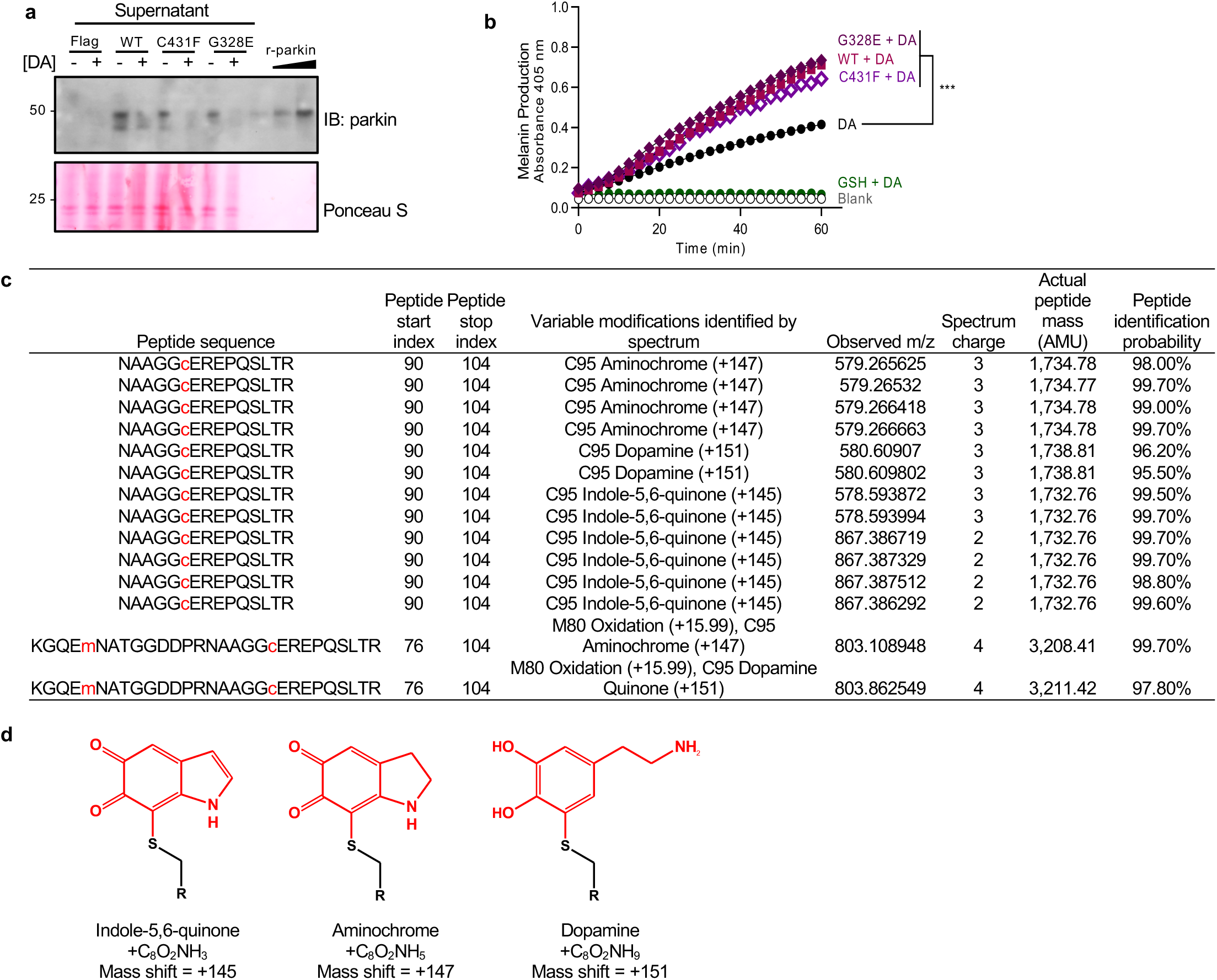
Parkin conjugates dopamine metabolites at C95. **(a)** Western blot analysis of ectopic, WT and mutant parkin expression in dopamine-treated M17 cells, as described in **Fig. 4a,b**. **(b)** *In vitro* melanin formation, as monitored by a colorimetric assay (absorbance rate, 405nm) demonstrating increased polymerization of dopamine (DA) over a 60 min period in the presence or absence (blank) of wild-type (WT) r-parkin and two point mutants (G328E; C431F). Melanin synthesis from dopamine is blocked in the presence of glutathione (GSH). Data represent the mean of triplicates ± SEM. ***p < 0.05 by 1-way ANOVA. **(c)** Table summarizing LC-MS/MS-based detection of adducts of dopamine-metabolites conjugated to C95 of r-parkin following exposure of r-parkin to aminochrome. Chemical structures for identified adducts are shown in **(d)**.

**Extended Data Figure 7.**
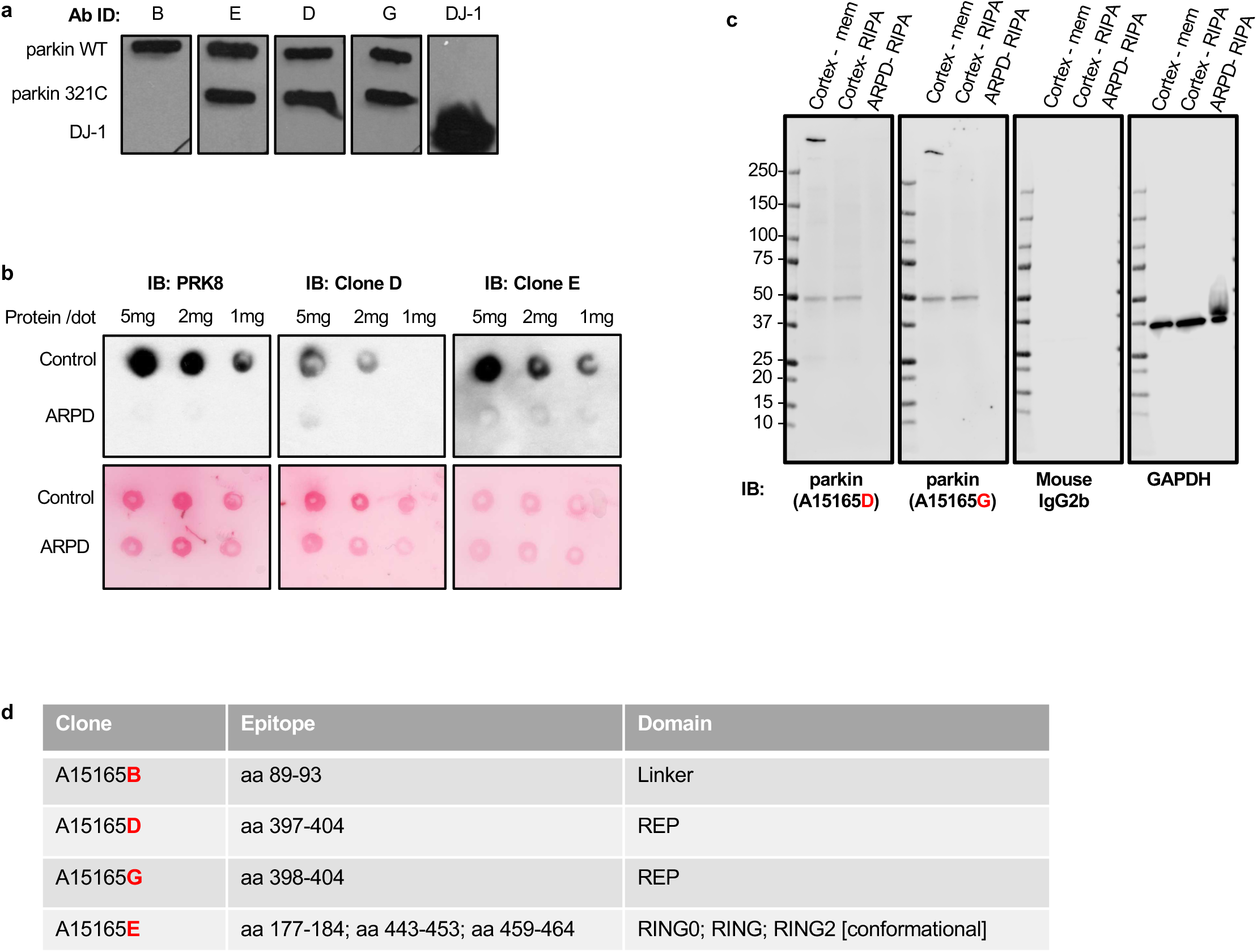
New monoclonal antibodies visualize parkin in ageing human midbrain. **(a-c)** Characterization of four murine, monoclonal antibodies (of IgG_2_ isotype; clone-B, -E, -D, and -G) in three different assays: **(a)** against recombinant (r-), full-length, untagged, wildtype (WT) human parkin using non-denaturing slot blots (100ng/slot) of original antigen as well as truncated r-parkin_321-465_ and full-length, untagged, human r-DJ-1; **(b)** against human brain lysates (SDS fractions from control and *PRKN-*linked ARPD cases) using non-denaturing dot blots; and **(c)** by denaturing SDS/PAGE and Western blotting of extracts from cortical specimens of a control brain and a parkin-deficient ARPD case. Screening by these three methods as well as by cell-based microscopy (using indirect immunofluorescence) revealed specific staining for four anti-parkin clones (-B, -E, -D and -G), which was conformation-dependent for clone-E. List of epitopes within the sequence of human parkin, as recognized by clones -B, -E, -D, and -G and informed by overlapping screening with 7-12 amino acid-long peptides covering full-length, human parkin. Note that the clone E epitope is conformational, comprised of the three regions indicated.

**Extended Data Figure 8.**
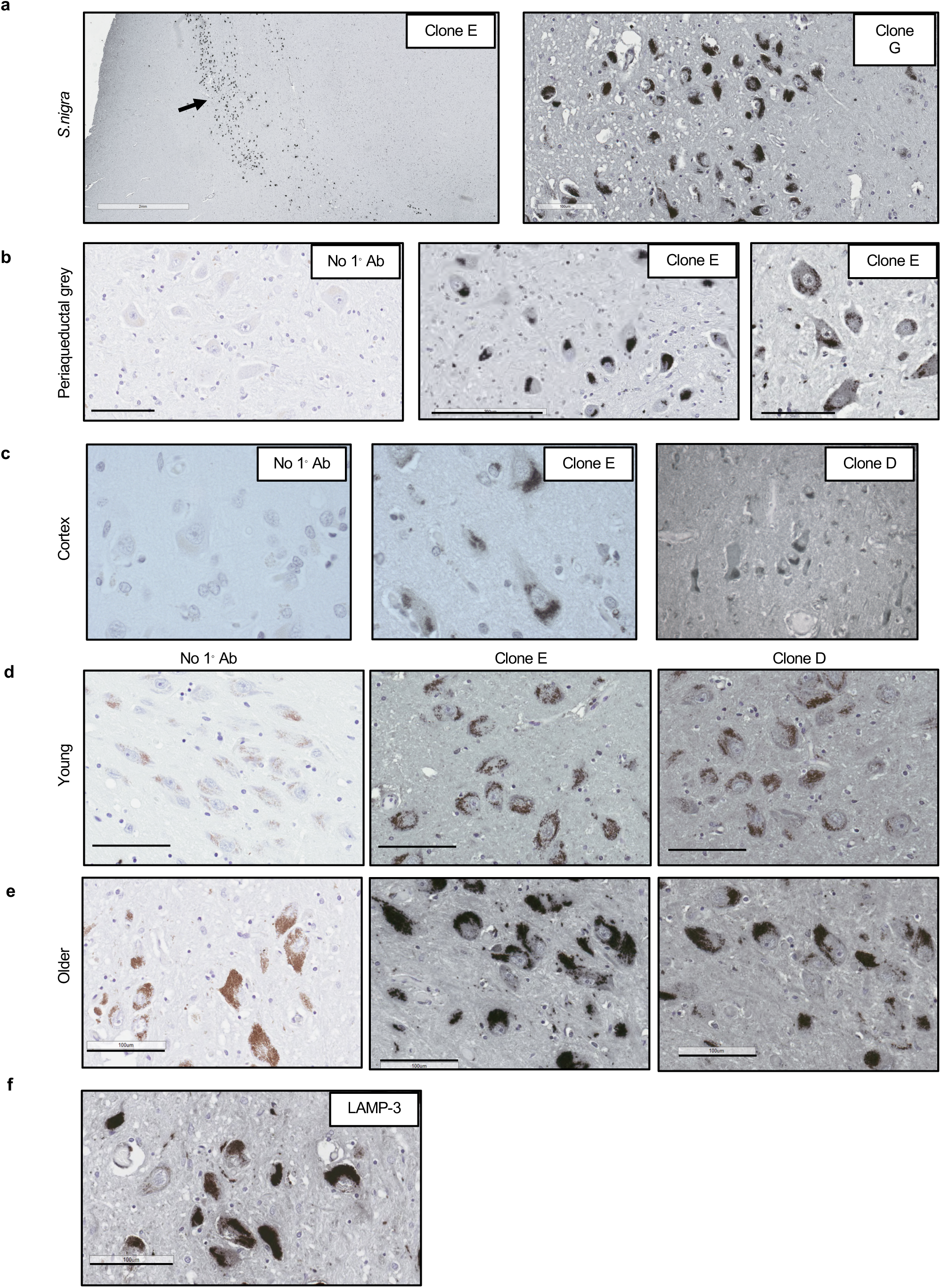
Parkin is readily detectable in human brain by routine microscopy. **(a-c)** Immunohistochemical detection of parkin in ageing human brain including **(a)** dopaminergic neurons of the S. nigra, **(b)** periaqueductal grey neurons in the midbrain and **(c)** cortex. Parkin was detected using the indicated clones and metal-enhanced DAB (black colour).. No primary Ab controls are shown. In all panels, scale bars represent 100 μm, or as indicated **(d-e)** Immunohistochemical detection of parkin in S.nigra tissue from from adult control subjects aged **(d)** 24 and **(e)** 66 years. Staining using clones E and D respectively and no primary antibody control are shown. Scale bars, 100 μm. **(f)** Immunohistochemical detection of LAMP-3 protein in dopaminergic neurons of the *S. nigra* from an adult control brain.

**Extended Data Figure 9.**
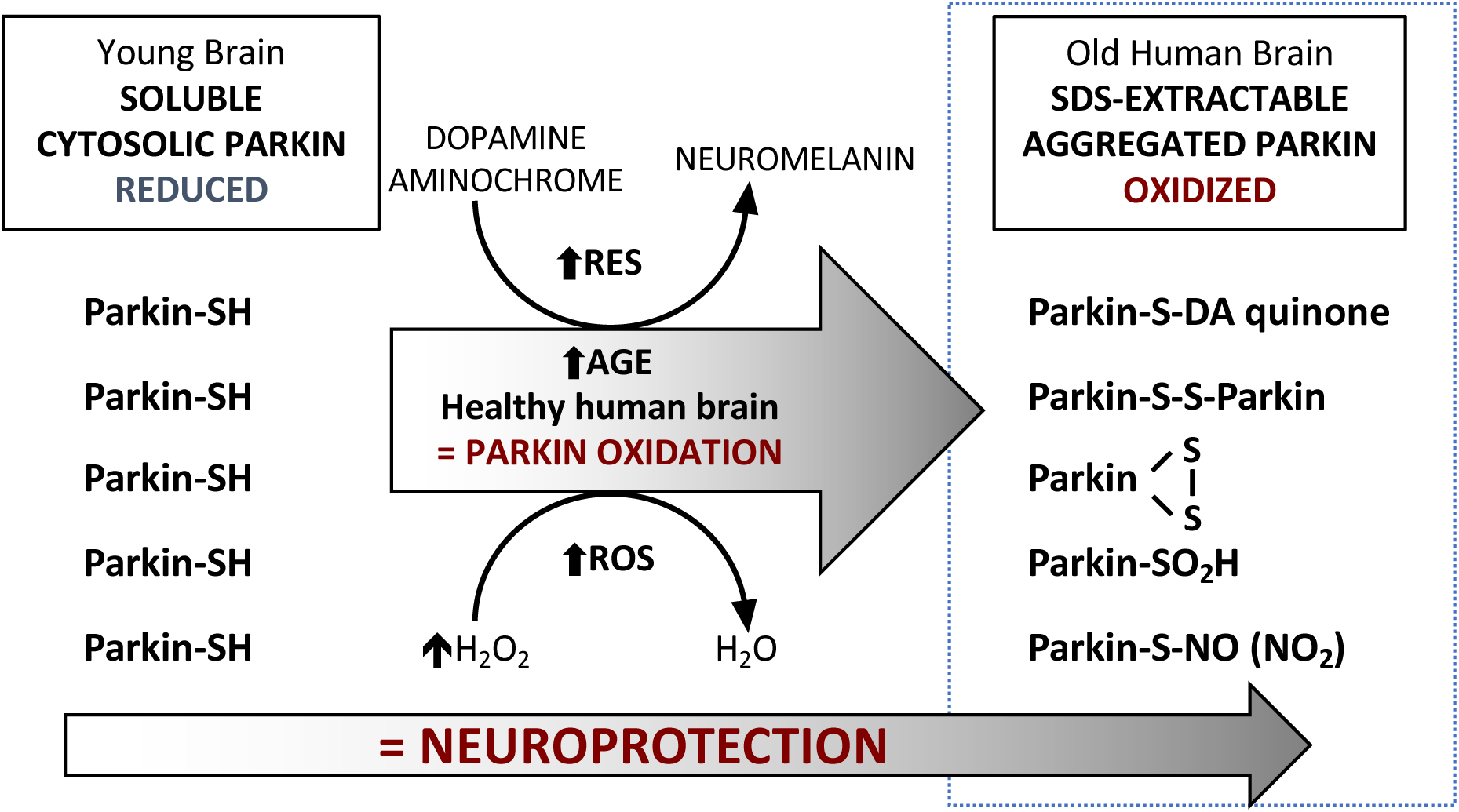
Proposed model for parkin’s redox functions in human midbrain. In human brain, parkin thiol (-SH) oxidation neutralizes cellular reactive oxygen species (ROS; H_2_O_2_) and potentially toxic dopamine (DA) metabolites (DA quinones; RES) during ageing. The oxidation of wild-type parkin promotes higher molecular weight species formation, insolubility and aggregation. During normal ageing, both reversible and irreversible oxidation events occur, which promote parkin’s eventual transition into an insoluble, oxidized state by the beginning of the 5th decade. The oxidation and aggregation lead to parkin’s accumulation within LAMP-3-positive lysosomes within DA neurons of the *S. nigra*. The multimodal oxidation of parkin confers neuroprotection. In *PRKN*-linked ARPD, the absence of parkin’s redox effects leads to a rise in ROS, reduced sequestration of DA radicals and elevation of RNS.

**Extended Data Table 1.** Summary of LC-MS/MS results with a focus on thiol redox state of parkin purified from human brain. Results for twelve IP runs (TS extracts; n=4; SDS extracts, n=8) of human cortices to identify reduced (IAA-tagged) and reversibly-oxidized (NEM-tagged) thiol residues. Parkin immunoprecipitation (IP) from human brains was performed using magnetic beads conjugated to the Prk8 antibody or a parkin monoclonal antibody (Clone B). Tissue extraction and IP was performed in the presence of IAA. Higher parkin coverage was consistently obtained from the TS-extracted fractions, along with a higher number of IAA-cysteines/total cysteines identified per run, when compared to the SDS-soluble fractions. In addition to IAA and NEM, we preselected for other non-reversible post-translation modifications for screening as indicated. In human brains, we detected irreversibly oxidized cysteines to sulfonic acid (trioxidation), at C95 and C253 and one NEM-labelled and oxidized cysteine, C253. Among the PTMs we preselected for, we also detected cysteine to dehydroalanine (Cys → Dha) shift and methionine oxidation.

|  | TS-extractable (IP, n=4)<br>Mean (range) | SDS-extractable (IP, n=8)<br>Mean (range) |
| --- | --- | --- |
| % Parkin protein coverage | 46.8 (35-72) | 19.4 (2-58) |
| # Peptides identified | 15.0 (10-23) | 6.4 (1-17) |
| # Cysteines identified | 17.5 (13-26) | 6.1 (1-16) |
| Reduced residues (IAA-Cys) | 17.5 (13-26) | 6.1 (1-16) |
| IAA/Total cysteines | 17.5/35 | 6.1/35 |
| % | 50 | 17.5 |
| IAA/Total identified cysteines | 17.5/28 | 6.1/28 |
| % | 62.5 | 21.8 |
| IAA/Total cysteines per run | 17.5/28 | 6.1/16 |
| % | 62.5 | 38.1 |
| Oxidized residues | Among PTMs preselected<br>for screening | Among PTMs preselected for<br>screening |
| Cys → Dha | Cys 95, 196, 238, 241, 253 | Cys 196 |
| NEM-cysteine | Not detected | Cys 253 |
| Sulfinic acid - Cys dioxidation | Not detected | Not detected |
| Sulfonic acid - Cys trioxidation | Cys 95, 253 | Not detected |
| Glutathionylation | Not detected | Not detected |
| S-nitrosylation | Not detected | Not detected |
| Dopamine quinone adduct<br>formation | Not detected | Not detected |
| Methionine oxidation | Met 80, 192, 327, 458 | Met 80, 192, 327, 458 |

## Footnotes

& El-Kodsi D et al., Parkinson Disease-Linked Parkin Lowers Cytosolic Oxidative Stress in Feedback Loop with Glutathione (2020), manuscript *in preparation*

