## Supplementary material for "Oxidative Modifications of Parkin Underlie its Selective Neuroprotection in Adult Human Brain": Tokarew et al_Supplementary Tables

| ID in figures | Sample ID | Brain region | Age | Sex | PMI (hrs) | Diagnosis | Parkin Solubility | H2O2/Tissue ratio |
|---|---|---|---|---|---|---|---|---|
| a | 1 | F ctx | 5 | F | 33 | HCO | 1 | |
| b | 2 | F ctx | 5 | F | 20 | HCO | 1 | 2.551 |
| | 3 | F ctx | 8 | M | 5 | HCO | 0 | 1.780 |
| t | 4 | F ctx | 13 | M | 13 | HCO | 1 | 2.217 |
| | 5 | F ctx | 15 | F | 9 | HCO | 1 | 1.349 |
| j | 6 | F ctx | 16 | F | 20 | HCO | 1 | |
| c | 7 | F ctx | 16 | F | 14 | HCO | 1 | 1.016 |
| ff | 8 | F ctx | 17 | M | 23 | HCO | 0 | 0.701 |
| | 9 | F ctx | 17 | M | 22 | HCO | 1 | |
| k | 10 | F ctx | 20 | F | 19 | HCO | 1 | |
| o | 11 | F ctx | 20 | M | 8 | HCO | 1 | |
| | 12 | F ctx | 20 | M | 6 | HCO | 0 | |
| | 13 | F ctx | 20 | M | 5 | HCO | 1 | |
| | 14 | F ctx | 21 | M | 30 | HCO | 1 | |
| | 15 | F ctx | 28 | M | 33 | NCO | 1 | |
| d | 16 | F ctx | 29 | F | 18 | NCO | 0 | 3.061 |
| | 17 | F ctx | 30 | M | 20 | HCO | 0 | 6.089 |
| | 18 | F ctx | 36 | M | 20 | HCO | 0 | 5.665 |
| u | 19 | F ctx | 37 | F | 13 | HCO | 0 | |
| v | 20 | F ctx | 38 | M | 17 | HCO | 0 | |
| | 21 | F ctx | 39 | M | 23 | HCO | 1 | |
| e | 22 | F ctx | 39 | M | 14 | HCO | 1 | |
| | 23 | F ctx | 42 | M | 18 | HCO | 1 | |
| f | 24 | F ctx | 43 | F | 22 | HCO | 0 | 4.601 |
| | 25 | F ctx | 44 | F | 21 | NCO | 0 | |
| | 26 | F ctx | 49 | F | 16 | HCO | 0 | 5.622 |
| s | 27 | F ctx | 49 | F | 14 | HCO | 0 | |
| m | 28 | F ctx | 54 | M | 16 | NCO | 0 | |
| | 29 | F ctx | 54 | F | 23 | NCO | 1 | |
| w | 30 | F ctx | 55 | F | 16 | HCO | 0 | 5.829 |
| q | 31 | F ctx | 56 | M | 23 | HCO | | |
| | 32 | F ctx | 57 | M | | HCO | 0 | |
| g | 33 | F ctx | 62 | M | 15 | NCO | 0 | 6.525 |
| x | 34 | F ctx | 65 | M | 5 | PSM | 0 | 6.473 |
| z | 35 | F ctx | 65 | M | 14 | PSM | 0 | 9.112 |
| n | 36 | F ctx | 65 | F | 42 | HCO | 0 | 5.768 |
| h | 37 | F ctx | 66 | M | | HCO | 0 | 6.674 |
| p | 38 | F ctx | 68 | M | 17 | HCO | 0 | 2.514 |
| i | 39 | F ctx | 70 | F | | HCO | 0 | 6.897 |
| r | 40 | F ctx | 70 | M | | HCO | 0 | 5.459 |
| aa | 41 | F ctx | 72 | M | | PSM | 1 | 6.274 |
| | 42 | F ctx | 75 | M | 48 | HCO | 1 | 2.878 |
| | 43 | F ctx | 75 | F | 13 | NCO | 0 | |
| y | 44 | F ctx | 75 | M | 17 | PSM | 0 | |
| | 45 | F ctx | 76 | M | 74 | PSM | 0 | |
| l | 46 | F ctx | 85 | F | 15 | NCO | 0 | |
| bb | 47 | Midbrain | 26 | M | 2 | NCO | 0 | |
| | 25 | Midbrain | 44 | F | 21 | HCO | 0 | |
| ee | 48 | Midbrain | 44 | F | 5 | NCO | 1 | |
| | 49 | Midbrain | 45 | M | 13 | NCO | 0 | |
| | 50 | Midbrain | 47 | F | 20 | NCO | 0 | |
| | 51 | Midbrain | 56 | M | 44 | PSM | 0 | |
| | 52 | Midbrain | 60 | M | 16 | PSM | 0 | |
| | 53 | Midbrain | 61 | M | 20 | NCO | 1 | |
| cc | 54 | Midbrain | 61 | M | 3.5 | NCO | 0 | |
| n | 36 | Midbrain | 65 | F | 42 | HCO | 0 | |
| dd | 55 | Midbrain | 65 | M | 6 | NCO | 0 | |
| | 41 | Midbrain | 72 | M | | PSM | 1 | |
| | 56 | Midbrain | 74 | F | | NCO | 0 | |
| | 42 | Midbrain | 75 | M | 48 | HCO | 0 | |
| | 57 | Midbrain | 75 | M | 70 | PSM | 0 | |
| | 45 | Midbrain | 76 | M | 74 | PSM | 0 | |
| | 58 | Midbrain | 79 | M | | PSM | 0 | |
| | 59 | Midbrain | 82 | M | 48 | PSM | 1 | |
| SC-1 | 60 | Spinal cord/muscle | 68 | F | 2.5 | NCO | 1 | |
| SC-2 | 61 | Spinal cord/muscle | 64 | F | 2 | HCO | 1 | |
| SC-3 | 62 | Spinal cord/muscle | 71 | | 2 | | 1 | |
| SC-4 | 63 | Spinal cord/muscle | 50 | M | 4 | | 1 | |

**Supplementary Table 1:** Summary of human brain tissue specimens used in this study with designated small letter identification, as used in the Figures. Abbreviations used: F ctx, frontal cortex; PMI, *post mortem* interval; hrs, hours; HCO, healthy control; NCO, neurological control; PSM, parkinsonism, n.d., not documented. For parkin solubility: 1, present in Tris-saline (TS) buffer; 0, not present in TS.

| | | r-parkin | | | | | | |
|---|---|---|---|---|---|---|---|---|
| | Treatment | Control | | $H_2O_2$ | | | | |
| | Run | IAA+NEM | IAA | 20µM | 1mM | 4.5mM | 4.5mM | 4.5mM |
| | Labelling | •IAA +NEM | | | | | | |
| Region | Cysteine Residues | | | | | | | |
| UBL | 59 | | • | n/d | • | • + | • | • |
| Linker | 95 | • | • | • | • | • + | • + | • + |
| RING0 | 150 | • | • | • | • | • + | • + | • |
| | 154 | • | • | n/d | • | • + | • + | • |
| | 166 | n/d | n/d | • | • + | n/d | • | • |
| | 169 | n/d | • | • | • + | n/d | • | • |
| | 182 | • | • | • | n/d | • + | • + | • |
| | 196 | • | • | • | • | • + | • + | • + |
| | 201 | • | • | • | • | • + | • + | • + |
| | 212 | • | • | • | n/d | • + | • + | • + |
| RING1 | 238 | • | • | • + | • | • + | • | • + |
| | 241 | • | • | • + | • | • + | • | • |
| | 253 | • | • | • + | • | • + | • | • |
| | 260 | • | • | n/d | n/d | • + | • | • |
| | 263 | • | • | n/d | n/d | • + | • | • |
| | 268 | • | • | n/d | n/d | • + | • | • |
| | 289 | • | • | n/d | n/d | • + | • + | • + |
| | 293 | • | • | n/d | • | • | • + | • + |
| | 323 | • | • | n/d | n/d | • + | • + | • + |
| IBR | 332 | • | • | n/d | n/d | • + | • | • |
| | 337 | • | • | n/d | • | • + | • | • + |
| | 352 | • | • | • | • | • | • + | • + |
| | 360 | • | • | • | • | • + | • | • |
| | 365 | • | • | • | n/d | • + | • | • |
| | 368 | • | • | n/d | n/d | • + | • + | • |
| | 377 | • | • | • | n/d | • + | • + | • + |
| RING2 | 418 | n/d | n/d | n/d | • | n/d | n/d | n/d |
| | 421 | n/d | n/d | n/d | • + | n/d | • + | • + |
| | 431 | n/d | • | n/d | • | n/d | • | • |
| | 436 | n/d | n/d | n/d | • + | n/d | • | • |
| | 441 | n/d | n/d | n/d | • + | n/d | • | • |
| | 446 | • | • | n/d | n/d | • + | • + | • + |
| | 449 | • | • | n/d | n/d | • + | • + | • |
| | 451 | • | • | n/d | n/d | • + | • | • |
| | 457 | • | • | • | • | • + | • | • + |

**Supplementary Table 2: Parkin's cysteine residues are redox active.**
Aliquots of human recombinant (r-) parkin that were oxidized by variable concentrations of $H_2O_2$ *vs.* control preparations were differentially labelled with iodoacetamide (IAA) and/or N-ethylmaleimide (NEM; as in Figure 4A) to identify reduced cysteines (IAA) or reversibly-oxidized residues (NEM). Proteins were subjected to LC-MS/MS and analyzed using Mascot Scaffold PTM to identify IAA (•) or NEM (+) adducts indicating when these were detectable on individual residues. Cysteines that were not detected as modified in individual runs are also listed (n/d). Note that cysteines within all four RING domains of parkin as well as in the linker and UbL domains can be variably modified. Corresponding data can be found in Extended Data Fig. 4h. For comparison, see **Extended Data Table 1** that lists modified cysteines identified in parkin purified from human brain.
